# Virus-assisted directed evolution of enhanced suppressor tRNAs in mammalian cells

**DOI:** 10.1101/2022.01.21.477302

**Authors:** Rachel E. Kelemen, Delilah Jewel, Rachel L. Huang, Zeyu Zhu, Xiaofu Cao, Muhammad Pasha, Jon Anthony, Tim van Opijnen, Abhishek Chatterjee

## Abstract

Site-specific incorporation of unnatural amino acids (Uaas) in living cells relies on engineered aminoacyl-tRNA synthetase/tRNA pairs borrowed from a distant domain of life. Such heterologous suppressor tRNAs often show poor intrinsic activity, presumably due to the failure to optimally interact with a non-native translation system. This limitation can be addressed in *E. coli* using directed evolution. However, no suitable selection system is currently available to do the same in mammalian cells. Here we report virus-assisted directed evolution of tRNAs (VADER) in mammalian cells, which employs a double-sieve selection scheme to facilitate single-step enrichment of active-yet-orthogonal tRNA mutants from naïve libraries. Using VADER, we developed improved mutants of *M. mazei* pyrrolysyl-tRNA, the most popular Uaa mutagenesis platform in eukaryotes. We also show that the higher activity of the most efficient mutants is specific for mammalian cells, alluding to an improved interaction with the unique mammalian translation system.

## Introduction

Site-specific incorporation of unnatural amino acids (Uaas) into proteins in mammalian cells holds much potential to enable both basic science as well as biotechnology applications.^1–4^ Central to this technology is a nonsense-suppressing aminoacyl-tRNA synthetase (aaRS)/tRNA pair which is engineered to charge the Uaa of interest without cross-reacting with any of its host counterparts. Such “orthogonal” aaRS/tRNA pairs are typically imported into the host cell from a different domain of life.^1–3^ The performance of the heterologous suppressor tRNA is often suboptimal in the new host, given that it must directly interact with a foreign translation system. Indeed, several studies have confirmed that Uaa incorporation efficiency in mammalian cells is limited by the poor performance of the heterologous suppressor tRNAs, which must be massively overexpressed to achieve acceptable levels of Uaa incorporation efficiency.^5–7^ The ability to overcome suboptimal tRNA performance will be beneficial to improve the robustness of the Uaa mutagenesis technology in mammalian cells, and to enable advanced applications.

The biology of tRNAs is complex and multifaceted (Supplementary Figure 1), involving expression, processing, post-transcriptional modifications, cellular stability, interaction with the cognate aaRS and the components of the translational system (e.g., elongation factors and the ribosome), etc.^8^ How these different facets of tRNA biology contribute to the poor performance of foreign suppressor tRNAs is poorly understood, which makes it challenging to develop better variants through rational design. However, directed evolution has been used with much success to develop improved orthogonal suppressor tRNA mutants for efficient Uaa incorporation in *E. coli*.^9–18^ This was made possible by the development of clever selection systems, which enables the enrichment of active and orthogonal suppressor tRNA mutants from large synthetic libraries. The ability to perform analogous tRNA evolution in mammalian cells has the potential to yield improved suppression systems, but no suitable platform is currently available. It is important to perform such directed evolution experiments in mammalian cells, instead of in lower organisms where directed evolution is better established, to ensure that the tRNA mutants are selected based on their improved interactions with the unique mammalian translation system.

Existing directed evolution strategies in mammalian cells largely rely on stable integration of the target gene in a cell line, followed by the creation of sequence diversity through untargeted or targeted random mutagenesis.^19–21^ This approach is unsuitable for tRNA evolution for two reasons: A) To ensure a clear genotype-phenotype connection, it is essential to have no more than a single genomically integrated mutant tRNA gene per cell. However, it has been shown that a large number of tRNA genes (>100) are needed per cell to achieve detectable Uaa incorporation efficiency;^5,7^ B) The low mutagenic frequency associated with these strategies is poorly suited for tRNA evolution, given its small size (<100 bp). One particular challenge arises when diversifying the stem regions of a tRNA, which are most frequently targeted for directed evolution: any mutation must be accompanied by a matching mutation on the other side to retain base-pairing, the loss of which compromises tRNA secondary structure and activity. Capturing such rich sequence diversity within the small tRNA gene is practically only feasible using synthetic site-saturation mutagenesis libraries. To enrich suppressor tRNA variants that are orthogonal and active in mammalian cells from such synthetic libraries, we need the following: i) controlled delivery of the library members into mammalian cells, such that each cell receives a single variant; ii) a selection scheme that enriches the active tRNA mutants, and removes crossreactive ones; and iii) the ability to identify the surviving mutants. We envisioned achieving these requirements by coupling the activity of the suppressor tRNA to the replication of a mammalian virus: i) encoding the library of tRNA variants in the virus genome would enable its controlled delivery to mammalian cells; ii) inserting a nonsense codon in an essential virus protein would render viral replication dependent on the activity of the suppressor tRNA, facilitating selective amplification of virions encoding active tRNA variants; and iii) the enriched tRNA sequences can be readily retrieved by isolating and sequencing the genome of the freshly amplified progeny virus. Some examples of virus-assisted directed evolution approaches in mammalian cells have been recently reported.^19,22,23^ However, these approaches use either natural or engineered error-prone replication of the virus genome to introduce sequence diversity, which is unsuitable for tRNA evolution due to the reasons described above. In addition, tRNA evolution demands simultaneous optimization of two distinct aspects: improving Uaa-incorporation activity, and suppressing potential cross-reactivity with the host aaRSs. No strategy currently exists to enable such sophisticated selection in mammalian cells.

We recently reported the feasibility of producing adeno-associated virus (AAV2) in mammalian cells, site-specifically incorporating Uaas into its capsid through amber suppression.^24,25^ In this system, successful AAV2 production is dependent on the activity of the suppressor aaRS/tRNA pair, providing an attractive platform for virus-assisted directed evolution of tRNA (VADER) in mammalian cells. AAV2 also offers additional advantages such as the lack of known pathogenicity, and a small genome that is amenable to facile manipulations.^26^ Here we report the development of an optimized double-sieve selection scheme (Figure 1a) for VADER that facilitates efficient one-step enrichment of active and orthogonal tRNA variants from naïve synthetic libraries. Using VADER, we developed mutants of *M. mazei* pyrrolysyl tRNA, which show significantly improved activity at lower expression levels. Furthermore, coupling VADER with next-generation sequencing (NGS) analysis provided a global view of the evolutionary landscape, where performance of each tRNA mutant in the entire library can be simultaneously tracked in mammalian cells from a single experiment. Finally, we show that the improved activities of the evolved tRNAs are host-specific, indicating that it likely stems from a better ‘fit’ with the unique translation apparatus of the mammalian cells.

**Figure 1.**
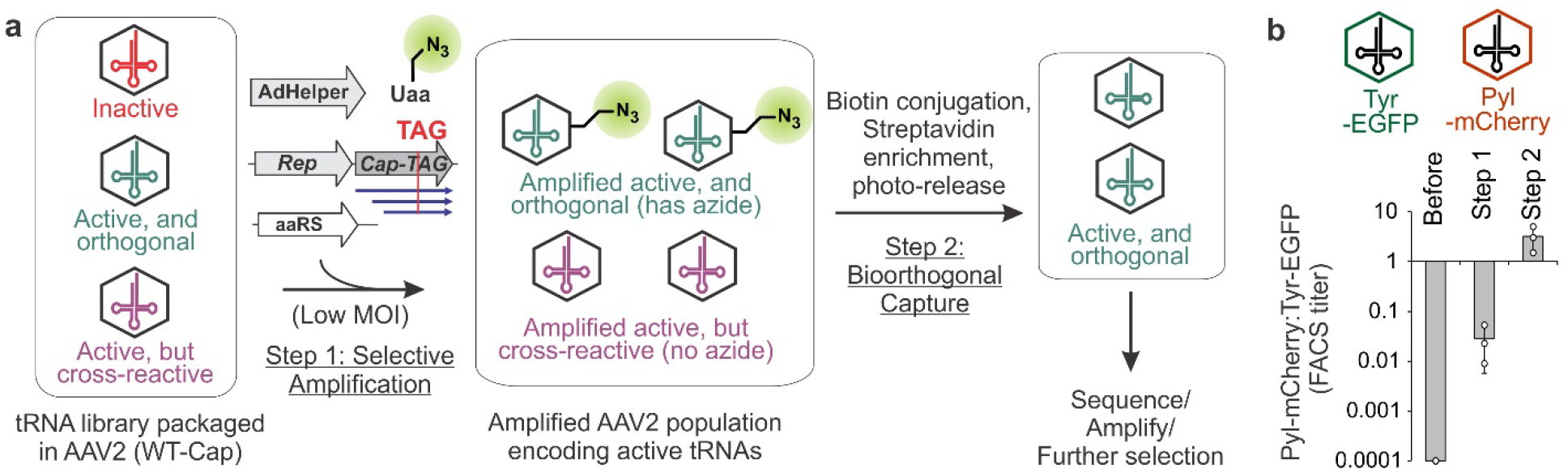
The VADER selection scheme. **a**, Mammalian cells are infected with AAV2 encoding the tRNA library at low MOI. Plasmids encoding TAG-mutant of Cap, other genetic components needed for AAV replication, and the cognate aaRS are provided *in trans* by transfection in the presence of a suitable azido-Uaa. Active and orthogonal tRNA mutants facilitate generation of packaged progeny AAV2 incorporating the Uaa into their capsid, which are isolated by chemoselective biotin conjugation followed by streptavidin pulldown. **b**, Two AAV2 vectors, encoding i) *E. coli* tRNA^Tyr^ and EGFP (Tyr-EGFP), and ii) tRNA^Pyl^ and mCherry (Pyl-mCherry), were mixed in a 10^4^:1 ratio and subjected to the VADER selection scheme using MbPylRS and its substrate AzK. FACS analysis of the surviving population shows a >30,000 fold cumulative enrichment of Pyl-mCherry. Data shown as mean ± s.d. (n = 3 independent experiments)

## Results and discussion

### Development and testing of the VADER selection system

Successful directed evolution relies on the delivery of no more than a single library member per cell. However, we have previously shown that the presence of a large number of tRNA genes per cell is required to observe detectable levels of nonsense suppression,^5–7^ posing a paradoxical challenge for developing VADER. We envisioned a solution to this challenge that takes advantage of the natural replication of the AAV2 genome in mammalian cells in the presence of its Rep gene (encoding multiple proteins that assist in viral replication) and additional helper genes from adenovirus (AdHelper), which would generate numerous copies of the incoming viral genome.^26^ We demonstrated that such replication is associated with a remarkable enhancement in transgene expression from the AAV2 genome by infecting HEK293T cells with an mCherry-encoding AAV2 in the presence or absence of Rep+AdHelper (provided *in trans*; Supplementary Figure 2).

Next, we evaluated if such amplification can produce sufficient levels of suppressor tRNA from a single incoming virion to support the expression of TAG-inactivated capsid gene and the production of progeny virus. HEK293T cells were infected with AAV2 encoding an mCherry reporter and an *M. mazei* pyrrolysyl-tRNA amber suppressor tRNA (tRNA_CUA_^Pyl^) at a low multiplicity of infection (MOI). Cells were then transfected with plasmids encoding the following components: i) Rep and Cap-454-TAG (the Cap gene of AAV2 encodes three overlapping capsid proteins, VP1-3; 454-TAG inactivates all three);^24,25,27^ ii) AdHelper genes; and iii) wild-type pyrrolysyl-tRNA synthetase from *M. barkeri* (MbPylRS). We observed progeny virus production only in the presence of the Uaa azido-lysine (AzK; Supplementary Figure 3), a substrate for MbPylRS. Efficiency of virus production was ~6% of an identical experiment, where wild-type Cap gene was used instead (Supplementary Figure 3). These experiments validate the basis of VADER: the ability to couple AAV2 replication to the activity of a suppressor tRNA encoded in its genome.

Synthetic tRNA libraries can harbor variants capable of cross-reacting with host aaRSs, so an additional strategy is necessary to remove such cross-reactive tRNA variants. We envisioned a novel approach to selectively enrich virions that encode active and orthogonal tRNAs based on their ability to incorporate a chemically unique Uaa into the AAV2 capsid, while the crossreactive ones would incorporate a canonical amino acid instead. If a Uaa with a bioorthogonal conjugation handle is used here, the resulting desired virus population can be isolated by chemoselective attachment of a biotin, followed by streptavidin enrichment (Figure 1a). We have previously demonstrated successful AzK incorporation and bioorthogonal modification of the AAV2 capsid.^25^ Using a photo-cleavable DBCO-biotin conjugate, we were indeed able to isolate AAV2-454-AzK following this strategy (Supplementary Figure 4). Together, the VADER selection scheme (Figure 1a) should enable the enrichment of an active and orthogonal tRNA population from a synthetic mutant library.

To show that an active suppressor tRNA can be enriched from a defined mixture of active and inactive tRNAs using VADER, we generated two AAV2 vectors; one encoding a tRNA_CUA_^Tyr^ (*E. coli* tyrosyl tRNA)^28^ and an EGFP, and another encoding tRNA_CUA_^Pyl^ and an mCherry (Figure 1b, Supplementary Figure 5). Packaged Pyl-mCherry and Tyr-EGFP virus were mixed in a 1:10^4^ ratio and subjected to the optimized VADER selection scheme (Figure 1a), using MbPylRS and its substrate AzK in step 1. MbPylRS charges AzK only to tRNA^Pyl^, and not tRNA^Tyr^;^29^ Pyl-mCherry virus should thus undergo enrichment relative to Tyr-EGFP. Indeed, FACS analyses revealed that the former is enriched approximately 280- and 130-fold after selective amplification and bioorthogonal capture steps, respectively, providing a >30,000-fold cumulative enrichment in a single round of selection (Figure 1b; Supplementary Figure 6).

### Application of VADER to generate tRNA^Pyl^ variants with improved activity

Next, we sought to use VADER for the directed evolution of tRNA^Pyl^, one of the most popular platforms for mammalian genetic code expansion (Figure 2a).^1–3,30^ Its acceptor (A) and T stems were chosen for diversification, as these regions typically interact with the components of the translation system.^31^ We created four different site-saturation mutant libraries covering both stems (Figure 2a), excluding the first base pair in the A-stem that is important for binding PylRS.^32^ The resulting libraries were cloned with >100 fold sequence coverage. Each library was subjected to the VADER scheme, and 30-50 surviving clones from each were characterized by sequencing. Base-pairing in these stem regions is important to maintain tRNA secondary structure and activity, but such fully base-paired sequences constitute a small fraction of the input tRNA library (Supplementary Figure 7). We were encouraged to find an enrichment of such fully base-paired sequences within the selected population, possibly indicating the successful selection of functional variants (Supplementary Figure 7; Supplementary Table 1, 2). Interestingly, sequenced selectants from the T-stem libraries revealed several wild-type or very similar mutants, while those from the A-stem libraries were more divergent (Supplementary Figure 7; Supplementary Table 1, 2). We evaluated the activity of each unique fully base-paired tRNA mutant isolated from the selection by co-transfecting them into HEK293T cells with MbPylRS and an EGFP-39TAG reporter in the presence and absence of 1 mM AzK. All tested mutants facilitated AzK-dependent expression of EGFP-39TAG, measured relative to the expression of a wild-type mCherry reporter encoded in the tRNA-plasmid (Figure 2d), confirming the selection of active and orthogonal mutants through VADER.

**Figure 2.**
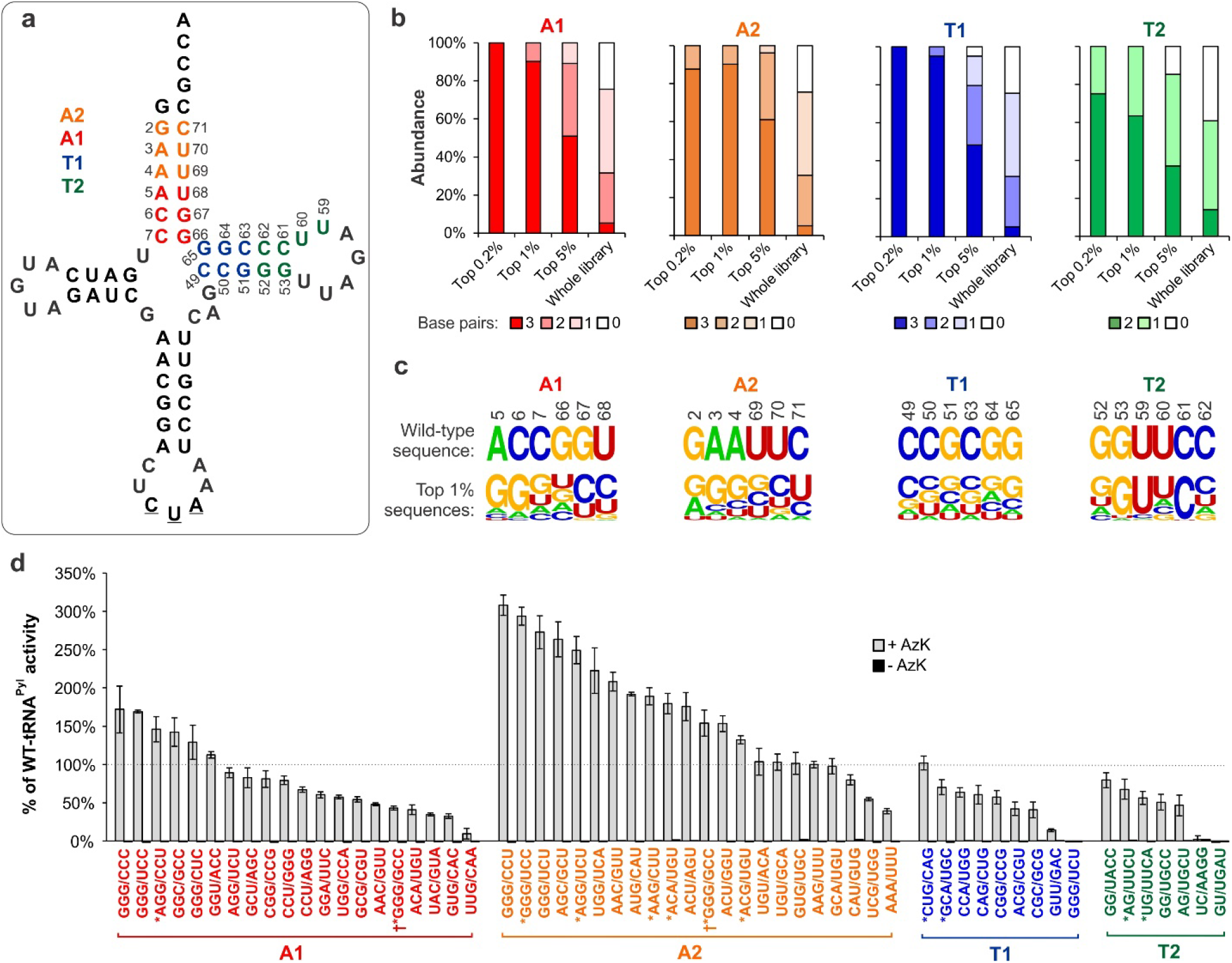
Directed evolution of tRNA^Pyl^. **a**, The sequences randomized to create four different libraries (A1, A2, T1, T2) of tRNA^Pyl^ are highlighted in four different colors. **b**, Degree of base pairing observed in the enriched mutants (through NGS analysis) upon the selection of each library. **c**, For each library, the wild-type sequence is shown on top and analysis of the 1% most-enriched sequences (n = 41) is shown below, revealing the relative abundance of each base at each randomized position. **d**, Efficiency of TAG suppression for unique tRNA^Pyl^ selectants measured using the EGFP-39TAG reporter. The tRNA encoded in the pAAV plasmid (also harboring a wild-type mCherry reporter) was co-transfected into HEK293T cells with MbPylRS and EGFP-39TAG in the presence or absence of 1 mM AzK. Expression of EGFP-39TAG was measured in cell-free extract, normalized relative to wild-type mCherry expression and plotted as a percentage of the normalized activity of wild-type tRNA^Pyl^. Data shown as mean ± s.d. (n ≥ 3 independent experiments). *sequences identified through NGS; †enriched sequences containing a G:G mispair.

To gain a more comprehensive look into the selection process, we used next-generation sequencing (NGS) to assess the composition of each of the four input libraries, as well as their selection output after VADER performed in duplicate. Gratifyingly, we were able to identify all possible mutants in each library and evaluate their relative abundance (Supplementary Figure 8). If a tRNA mutant is more active, it would facilitate higher Cap-TAG expression, and the production of more progeny virus. Consequently, the degree of enrichment of each tRNA mutant upon selection, which can be calculated from its normalized abundance before and after the selection (by NGS; Supplementary Figure 9), should correlate with its activity in mammalian cells. Indeed, we found that the strongly enriched mutants (top 0.2% – 1%) from each library were heavily base-paired (Figure 2b), a hallmark of functional tRNAs. Additionally, for the individually characterized tRNA mutants (Figure 2d), we found a strong correlation between their relative enrichment and the observed suppression efficiency (Supplementary Figure 10), further confirming that the degree of enrichment factor upon selection is a reliable indicator of tRNA performance in mammalian cells. We identified several highly enriched sequences from each library that were not identified by sequencing individual surviving clones. We synthesized several such mutants and characterized their activity relative to the wild-type tRNA^Pyl^ (indicated by a * in Figure 2d). Cumulative characterization of all sequences revealed many A-stem mutants with significantly higher activity than the wild-type tRNA^Pyl^ (Figure 2d), with the most active variant (tRNA^Pyl^-A2.1) showing a 3-fold improvement. In contrast, the T-stem libraries yielded fewer active variants (Supplementary Figure 9), and none showed higher activity (Figure 2d).

NGS-coupled VADER provides a global view of the selection process, where the performance of each mutant in mammalian cells can be tracked across the entire library (Supplementary Figure 9). In addition to identifying individual top-performers, it revealed many additional nuances. For example, mispairing within tRNA stem regions is generally detrimental for its function, and such sequences identified from a traditional selection output would be typically ignored as artifacts. However, we identified sequences containing G:G mispairs in both A-stem libraries that were highly enriched upon selection, suggesting that these are legitimate hits. We tested two such mutants (indicated with a † in Figure 2d), both of which were active and orthogonal; the sequence GGG/GCC from A2 library actually showed 50% higher activity than WT-tRNA^Pyl^. Additionally, the global enrichment profile of each library (Supplementary Figure 9) revealed that a large fraction of the A-stem libraries showed significant enrichment, indicating remarkable sequence plasticity of this region. In contrast, a much smaller fraction of the T-stem libraries showed significant enrichment, highlighting its limited tolerance for sequence alterations. Finally, analyses of all highly enriched (top 1%) sequences from the NGS-coupled VADER revealed interesting trends (Figure 2c). For example, most enriched mutants from the A-stem libraries show a sequence preference that is distinct from the wild-type tRNA^Pyl^: these are significantly more G:C rich, with a strong preference for G residues at positions 2-7, and pyrimidines on the 3’ side of the A-stem. Characterization of individual clones corroborates that these sequence features are associated with enhanced activity. In contrast, the best-performing mutants from the T-stem libraries largely mirrored the wild-type sequence. Poor understanding of foreign tRNA biology in mammalian cells currently limits our ability to shed light on the mechanistic underpinnings of these observations. However, high-throughput characterization of large tRNA mutant libraries in this way can create an empirical knowledge base that will aid the design of improved variants.

### Further characterization of tRNA^Pyl^-A2.1

More active tRNA^Pyl^ mutants identified through VADER can improve the robustness and scope of the Uaa mutagenesis technology in mammalian cells using the popular pyrrolysyl platform. To this end, we further benchmarked the performance of the most efficient mutant, tRNA^Pyl^-A2.1, against its WT counterpart. Even though this mutant was selected on the basis of incorporating AzK into the AAV2 capsid, we expected it to maintain its enhanced activity for other Uaas genetically encoded using the pyrrolysyl pair. Indeed, in addition to AzK (Figure 3c), tRNA^Pyl^-A2.1 also facilitated improved incorporation of N^ε^-acetyllysine (AcK; Figure 3b, d) using a previously established mutant MbPylRS.^33^ Furthermore, A2.1 facilitated incorporation of AcK – a weak substrate – into two different sites in the reporter, whereas the expression of the same reporter using WT tRNA^Pyl^ was near background level (Figure 3d). We also constructed opal and ochre (UGA and UAA) suppressor variants of tRNA^Pyl^-A2.1, each of which exhibited significantly higher activity relative to WT-tRNA^Pyl^ (Figure 3e). In fact, the opal suppression efficiency of A2.1 was on par with that of amber suppression using WT-tRNA^Pyl^ (Figure 3c, 3e). This will be particularly beneficial for site-specific incorporation of two distinct Uaas, which requires suppression of two different nonsense codons, and has been limited by the poor efficiency of non-UAG nonsense codons by the WT-tRNA^Pyl^.^29,34^ We further validated the enhanced activity of tRNA^Pyl^-A2.1 by incorporating AzK into the full-length human epidermal growth factor receptor (EGFR) at a significantly higher efficiency (Figure 3f).

**Figure 3.**
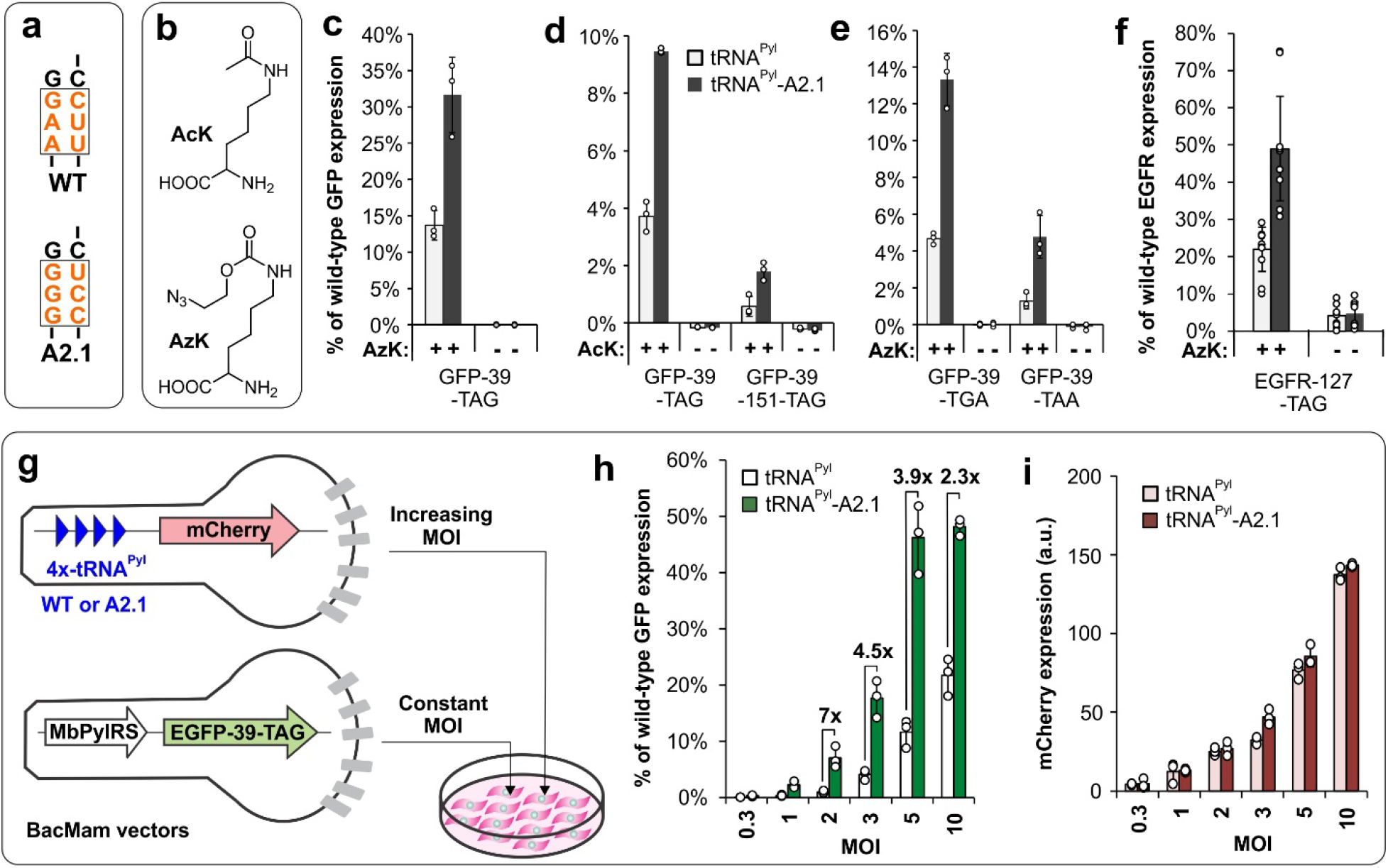
Characterization of tRNA^Pyl^-A2.1 activity. **a**, Sequences of wild-type (WT) and A2.1. **b**, Structures of AcK and AzK. **c**, Expression of EGFP-39TAG using the WT or A2.1 tRNA with MbPylRS, in the presence (+) and absence (-) of AzK. **d**, Expression of EGFP-39TAG and EGFP-39TAG-151TAG using the WT or A2.1 tRNA with AcK-selective MbPylRS, in the presence and absence of AcK. **e**, Expression of EGFP-39TGA and EGFP-39TAA using tRNA_UCA_^Pyl^ and tRNA_UUA_^Pyl^ (for both WT and A2.1), respectively, and MbPylRS in the presence and absence of AzK. **f**, Expression of EGFR-127TAG (with a C-terminal EGFP fusion) using the WT or A2.1 tRNA with MbPylRS, in the presence and absence of AzK. **g**, Mammalian cell-optimized baculovirus (BacMam) vectors and the scheme of the experiment for **h**-**i**. **h**, Expression of EGFP-39TAG as increasing MOI of tRNA-BacMam vector is used. **i**, Expression of mCherry in the same experiments as **h** shows equivalent delivery of both tRNAs, which increases proportionately with higher MOI. Expression of the fluorescent reporters is measured in HEK293T cell-free extract. Also see Supplementary Figure 11 for the corresponding images. For **c**-**f** and **h**, EGFP-39TAG expression is reported relative to wild-type EGFP expression from the same vector. Data shown as mean ± s.d. (n ≥ 3 independent experiments).

So far, we tested the activities of tRNA^Pyl^ variants by transiently transfecting the appropriate plasmids. However, transient transfection delivers a large number of plasmids into cells, resulting in an overexpression of the encoded genes.^7^ We have shown in the past that such overexpression can confound the comparison between two systems in mammalian cells: the performance of the more active counterpart can get saturated at higher expression, allowing the weaker variant to ‘catch up’.^7^ To compare the activity across varying expression levels, we previously developed a mammalian-optimized baculoviral (BacMam) delivery vector, which enables convenient tuning of transgene expression by simply altering the virus-to-cell ratio.^7^ This controlled delivery system was used to further compare the activity of tRNA^Pyl^-A2.1 to its WT counterpart. We used two BacMam vectors in this assay: one that encodes MbPylRS and EGFP-39TAG, and another encoding four copies of the tRNA^Pyl^ (either wild-type or A2.1) as well as a wild-type mCherry reporter (Figure 3g). The former was used at a constant MOI, while the latter was gradually increased from an MOI of 0.3 to 10 (Figure 3g). The mCherry and the EGFP signals represent the amount of tRNA-encoding vectors delivered, and the resulting UAG-suppression activity, respectively. The EGFP signal was normalized relative to cells infected with a separate BacMam vector encoding a WT-EGFP reporter at the same MOI. The mCherry expression levels increased linearly with increasing amount of tRNA-vector used, and signals from both tRNA vectors were comparable (Figure 3i, Supplementary Figure 11). When the tRNA-encoding vectors were delivered at low MOIs of 1 and 2, EGFP-39TAG expression (relative to WT-EGFP) facilitated by A2.1 was 2% and 7%, respectively, whereas the same for WT-tRNA was undetectable and 1% (Figure 3h, Supplementary Figure 11). Above MOI 5, A2.1 activity became saturated at ~50% reporter expression, while the WT-tRNA activity continued to close the efficiency-gap. These observations provide a more nuanced view of their relative activities, where A2.1 shows significantly higher benefit under copy-number limited scenarios. This property is expected to be particularly beneficial for advanced applications of the Uaa technology, where there are practical limits on the number of tRNA copies that can be delivered (e.g., *in vivo* applications,^35^ development of stable cell lines,^5^ etc.).

### The origin of the improved activity of tRNA^Pyl^-A2.1

As discussed earlier, the connection between the poor performance of foreign tRNAs and various facets of tRNA biology remains unclear. It also limits our ability to comprehend the molecular basis of the improvement achieved through directed evolution. Indeed, many orthogonal tRNAs have been evolved in *E. coli*, but the underlying mechanism for their improved activity is not understood.^9–18^ A mutant tRNA might exhibit higher activity simply due to higher cellular abundance (e.g., due to higher stability). We used Northern blot analysis to show that, upon transient transfection of HEK293T cells, A2.1 is actually expressed at a lower level relative to WT-tRNA^Pyl^ (Figure 4a), suggesting that it is intrinsically more active.

**Figure 4.**
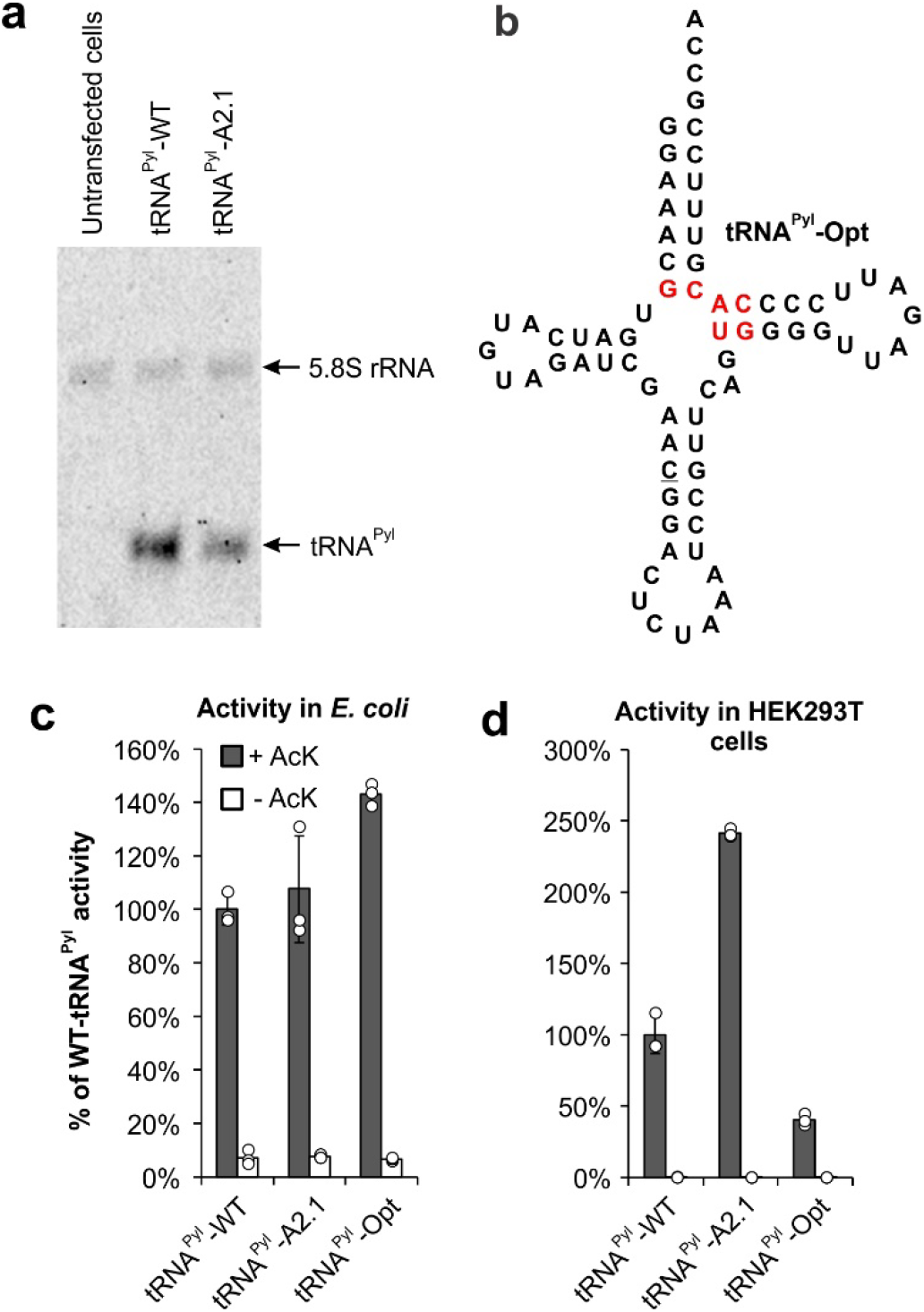
Origin of the improved activity of tRNA^Pyl^-A2.1. **a**, Northern blot analysis shows lower expression of A2.1 relative to WT, when each tRNA^Pyl^ has been transfected into HEK293T cells. **b**, Sequence of tRNA^Pyl^-Opt (optimized in *E. coli)* showing key mutations in red. **c**, Activity of tRNA^Pyl^-WT, A2.1, and Opt in *E. coli* measured using the sfGFP-151-TAG reporter. **d**, Activity of tRNA^Pyl^-WT, A2.1, and Opt in HEK293T cells measured using the EGFP-39-TAG reporter. For both c and d, a mutant MbPylRS selective for AcK was used and the activities were measured in the presence and the absence of AcK. Expression of the fluorescent reporter was measured in cell-free extract and reported as the % of WT-tRNA^Pyl^ activity. Data shown as mean ± s.d. (n = 3 independent experiments).

Due to its universal orthogonality, the pyrrolysyl pair can be used for Uaa mutagenesis in both *E. coli* and mammalian cells. A tRNA^Pyl^ mutant (tRNA^Pyl^-Opt; Figure 4b) was previously developed in *E. coli* that shows modest improvement in activity relative to tRNA^Pyl^-WT.^36,37^ We wondered how tRNA^Pyl^-A2.1 and tRNA^Pyl^-Opt, which have been engineered for higher activity in mammalian cells and *E. coli*, respectively, would perform in a different host cell. If the higher activity engineered can be transferred to different host, it would indicate that a factor intrinsic to this pair (e.g., PylRS-tRNA^Pyl^ interaction or tRNA stability) underlies the improvement. Alternatively, if the improvement is specific for a particular host, this would suggest that an improved interaction with the translation system of the particular host leads to the higher activity. We compared the activities of tRNA^Pyl^ WT, A2.1 and Opt in both *E. coli* and HEK293T cells using the same MbPylRS mutant (which charges AcK).^33^ As expected, A2.1 and Opt showed higher activity than WT in HEK293T cells (Figure 4d), and *E. coli* (Figure 4c), respectively. However, the activity of A2.1 in *E. coli* was comparable with WT, while Opt showed a significantly weaker activity in mammalian cells relative to WT. These observations strongly suggest that the enhanced activity of A2.1 originates from an improved interaction with the unique translation system of mammalian cells. It also underscores the importance of performing tRNA evolution directly in the target host to ensure proper ‘fit’ with its unique translation machinery. This is in contrast with the paradigm in aaRS engineering, where enhanced mutants developed in *E. coli* are routinely imported into mammalian cells with retention of function.^28,38,39^

## Conclusion

In summary, we have developed a general strategy to systematically address the poor performance of suppressor tRNAs in mammalian cells using a novel virus-assisted directed evolution platform. Coupling VADER with NGS facilitated a global analysis of how each member of our synthetic tRNA mutant libraries perform in mammalian cells, revealing sequence traits that positively or negatively affect their performance. Application of this technology to synthetic libraries of tRNA^Pyl^ revealed a high degree of evolvability of its A-stem, but not the T-stem, and led to the identification of many A-stem mutants with improved activities. The most active mutant (A2.1) was found to express at a lower level relative to tRNA^Pyl^-WT, pointing at its higher intrinsic activity. We also established an assay to compare the activities of different tRNA mutants across a range of different expression levels by controlled delivery of a defined copies using a baculovirus vector. This analysis revealed a significantly higher benefit of A2.1 at lower expression levels. Finally, we showed that enhanced tRNA^Pyl^ activity achieved through directed evolution is host-specific, suggesting that a better interaction with the non-native host translational apparatus likely underlies their improved performance. It should be possible to extend the application of VADER to other suppressor tRNAs commonly used for genetic code expansion in mammalian cells. Additionally, this platform can be adapted for the directed evolution of other biological machinery in mammalian cells for Uaa mutagenesis and beyond.

## Acknowledgements

We thank the National Science Foundation (MCB-1817893), and National Institutes of Health (R35GM136437 to A.C. and U01 AI124302 to T.v.O.) for financial support.

## Author contribution

A.C. and R.E.K. designed the project, R.E.K. developed and optimized the VADER system and performed selections, D.J. performed additional selections and characterized the tRNA mutants, M.P. and X.C. assisted in cloning, T.v.O, Z.Z and J.S.A were involved in Illumina sequencing, design and analyses, and A.C., R.E.K., and D.J. prepared the manuscript.

### Competing interests

A patent application has been submitted on the improved tRNA mutants reported here. A.C. is a senior advisor at BrickBio, Inc.

### Additional information

Supplementary information is available for this paper.

## Notes

### Competing Interest Statement

AC is a cofounder and senior advisor at BrickBio, Inc.

