## Supporting information for "Virus-assisted directed evolution of enhanced suppressor tRNAs in mammalian cells"

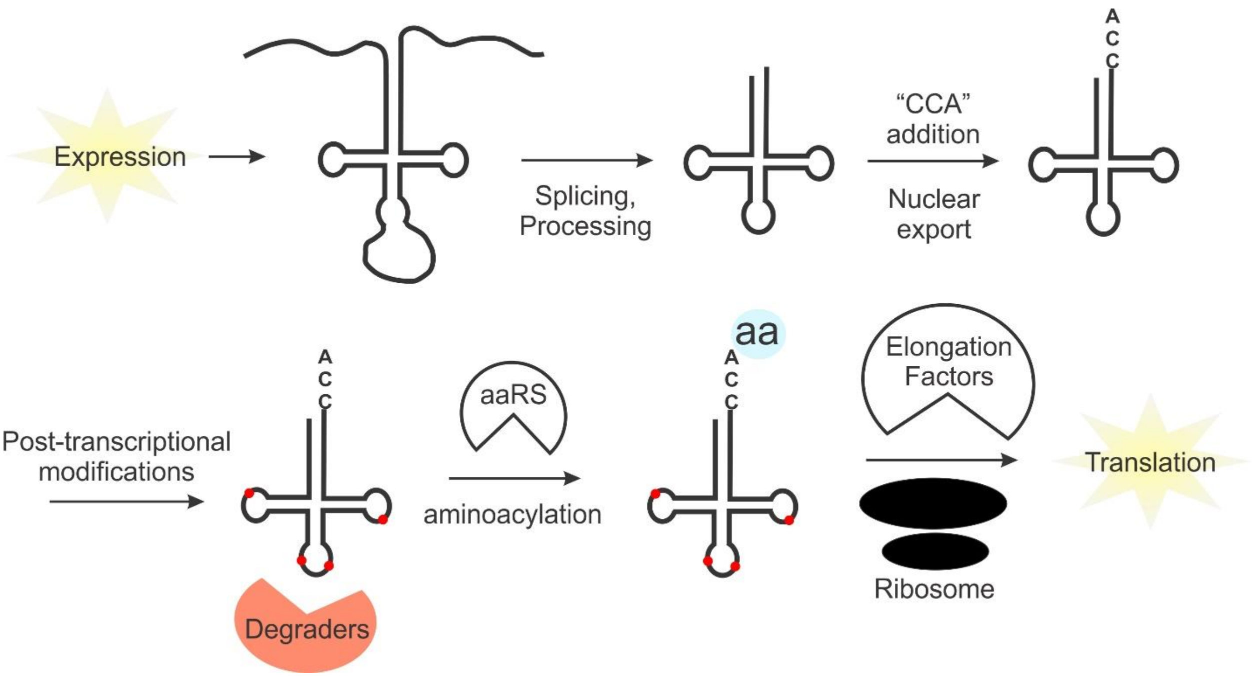

**Supplementary Figure 1**: The tRNA biology is complex and multifaceted. For proper processing and function, a foreign tRNA must rely on direct interactions with numerous host machinery. Suboptimal nature of these interactions – which have not been optimized through evolution – can compromise the intrinsic efficiency of foreign tRNAs. The major factors contributing to the poor efficiency of foreign tRNAs in mammalian cells remain poorly understood.

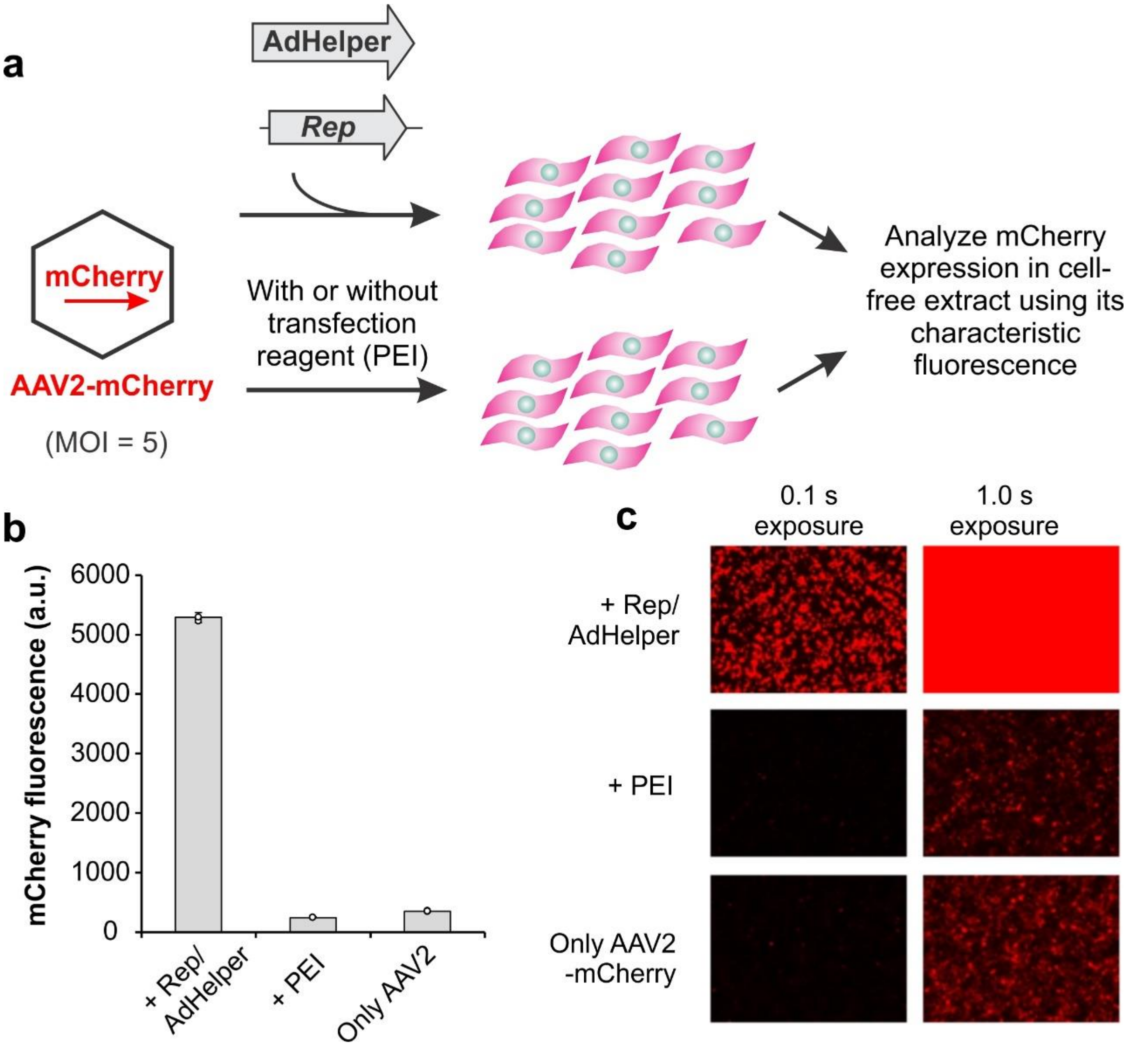

**Supplementary Figure 2**: Replication of the AAV2 genome in mammalian cells is associated with a large increase in transgene expression. **a**, Scheme of the experiment. HEK293T cells were infected with AAV2 encoding a wild-type mCherry gene, in the presence or absence of co-transfection of plasmids harboring Rep and AdHelper genes. Expression of mCherry was monitored 48 hours post-transfection by fluorescence microscopy (**c**), as well as by measuring its characteristic fluorescence in cell-free extract (**b**). The mCherry expression is slightly diminished in the presence of the transfection reagent (PEI) alone, possibly due to a perturbation of viral entry. Data shown as mean ± s.d. (n = 3 independent experiments).

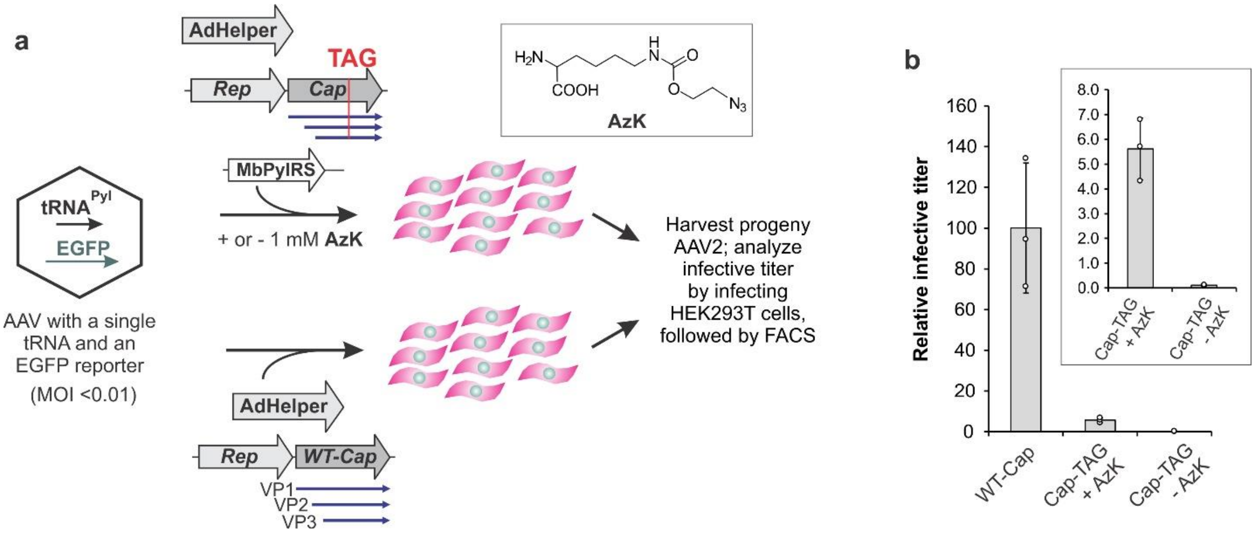

**Supplementary Figure 3**: A single AAV2-encoded tRNA gene can facilitate the expression of TAG-inactivated capsid gene (Cap) and the production of progeny virus. **a**, Scheme of the experiment. HEK293T cells are infected with AAV2 encoding a $tRNA_{CUA}^{Pyl}$ and a wild-type EGFP gene at a low MOI, then further transfected with plasmids encoding: i) AAV2 Rep and Cap-454-TAG genes, ii) MbPylRS, and iii) AdHelper in the presence or absence of 1 mM AzK. The feasibility of packaging AAV2 incorporating AzK at the 454 position of Cap, and that it does not perturb the virus, have been previously demonstrated.[1] Suppression of the TAG codon at 454 position of Cap leads to AzK incorporation into all three overlapping capsid proteins, VP1, VP2 and VP3 (60 total copies), at a surface exposed site. An identical experiment in which Cap-454-TAG is replaced by a wild-type Cap was also performed. After 48 hours, the progeny virus was harvested from these cells and titered by infecting freshly seeded HEK293T cells, followed by FACS analysis. **b**, AzK-dependent production of progeny virus is observed when Cap-454-TAG is used (magnified in the inset); the efficiency is significantly lower than the identical experiment where wild-type Cap is used instead. Data shown as mean ± s.d. (n = 3 independent experiments).

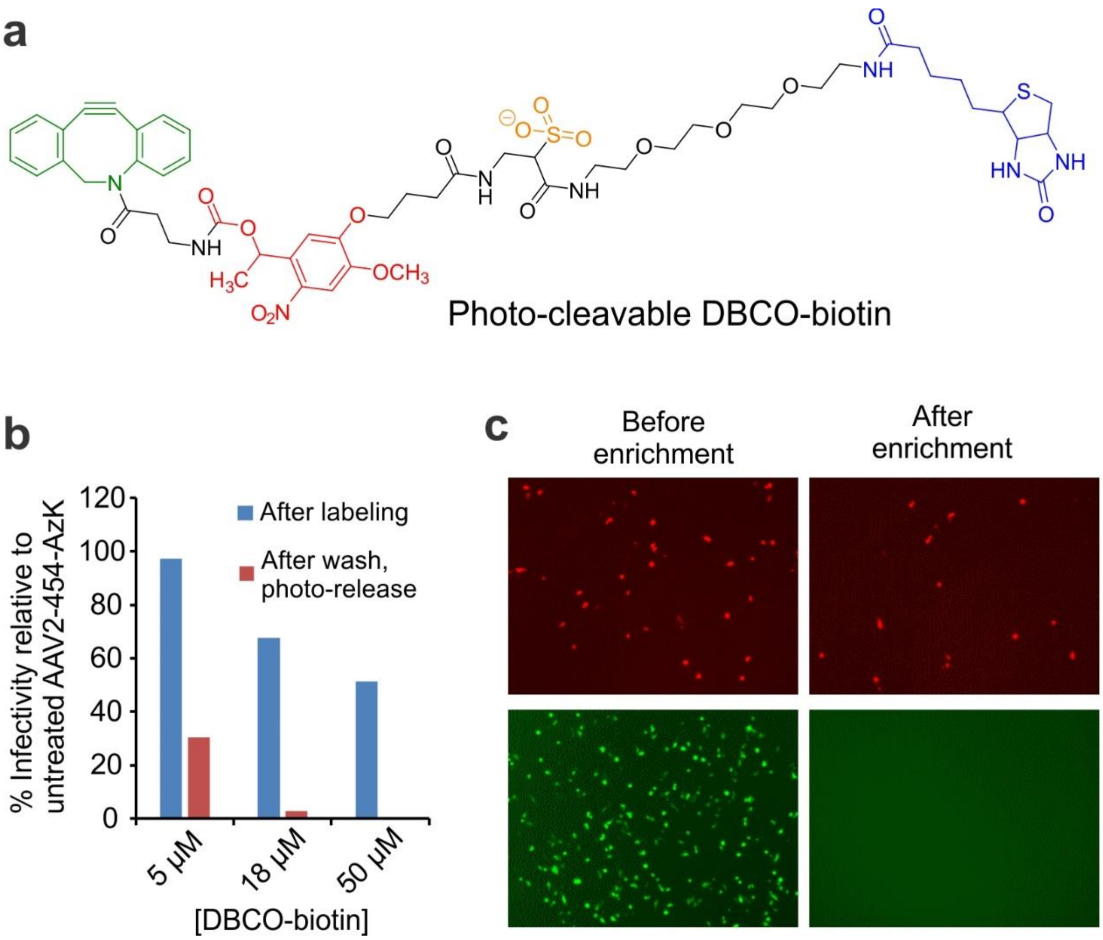

**Supplementary Figure 4**: AAV2-454-AzK can be isolated by bioorthogonal attachment of a photo-cleavable DBCO-biotin conjugate followed by streptavidin binding and photorelease. **a**, Structure of the photocleavable DBCO-biotin conjugate. **b**, AAV2-454-AzK was treated with different concentrations of the DBCO-biotin for 1 hr, the reaction was quenched using excess AzK, and unreacted small molecules were removed by dialysis. The biotin-labeled virus was captured using streptavidin-agarose, then released by 365 nm irradiation. The infective titer of the virus was measured (by infecting HEK293T cells followed by FACS) before treatment, after biotin modification, and after photo-release, then normalized for volume change. Infectivity relative to untreated virus was plotted. We find that a low degree of biotinylation (at 5 µM reagent; each virion harbors 60 AzK residues) does not affect the AAV2 infectivity, but increased modification of the capsid (at higher DBCO-biotin concentrations) does. Also, using 5 µM DBCO-biotin, modified virus is recovered from streptavidin resin with good efficiency (~30%), whereas the yield is poor when a higher reagent concentration is used. **c**, Using this optimized labeling/capture strategy, AAV2-454-AzK encoding an mCherry reporter can be enriched from its mixture with an EGFP-encoding AAV2 with wild-type capsid (no azide). Fluorescence microscopy images of HEK293T cells infected with the mixed virus population before and after the selection are shown.

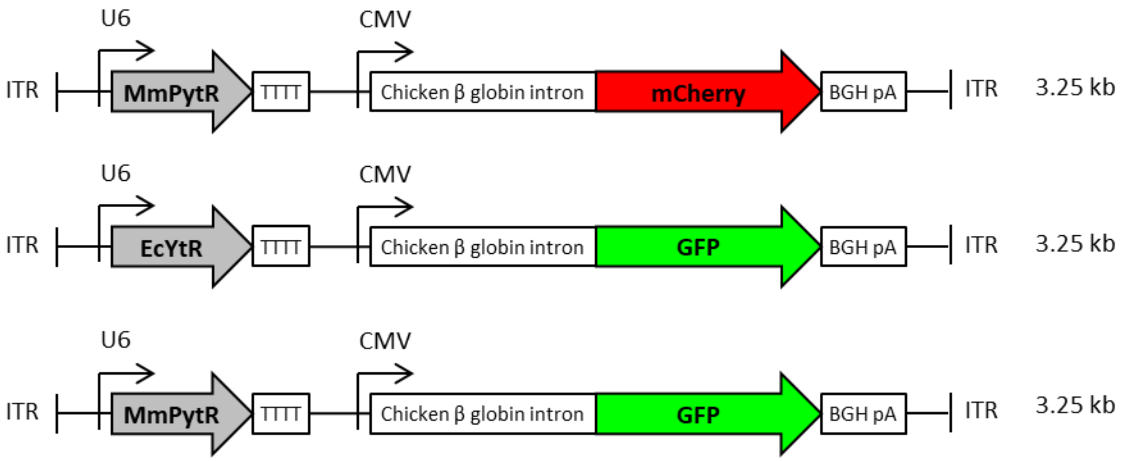

**Supplementary Figure 5.** Schematic maps of AAV2 cargoes containing various tRNAs and fluorescent proteins used in this study.

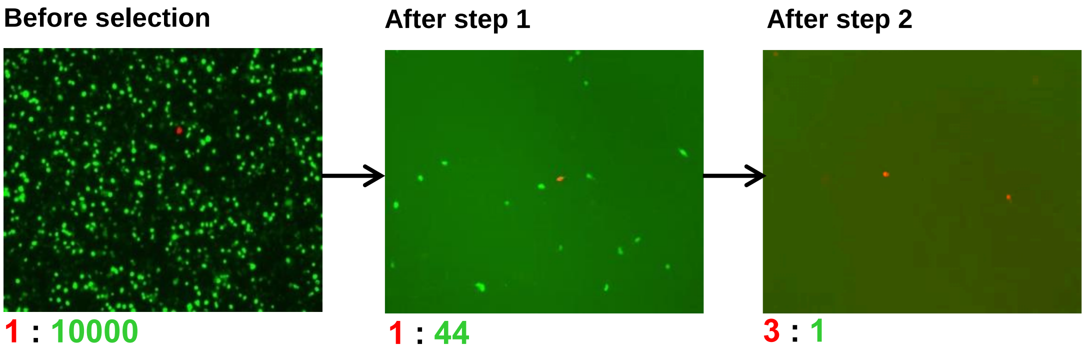

**Supplementary Figure 6**. Representative microscopy images of cells infected with the mixed virus population (Tyr-EGFP : Pyl-mCherry) before selection, after step 1, and step 2. Merged images from the EGFP and mCherry channels are shown for each. The ratios of the two viruses (Pyl-mCherry : Tyr-EGFP) as measured by FACS for these experiments are shown below the images.

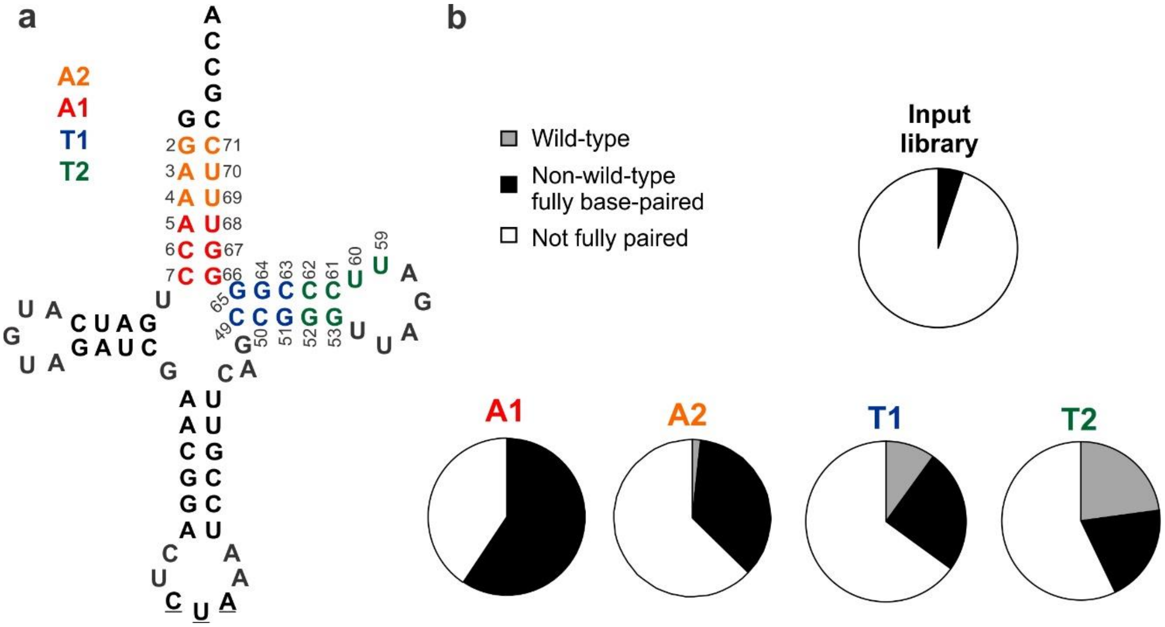

**Supplementary Figure 7**. Analysis of tRNA sequences emerging from the selection of each library (see Supplementary Table 2). The theoretical abundance of fully base-paired, not fully base-paired and the wild-type sequence in the input library is also shown for reference.

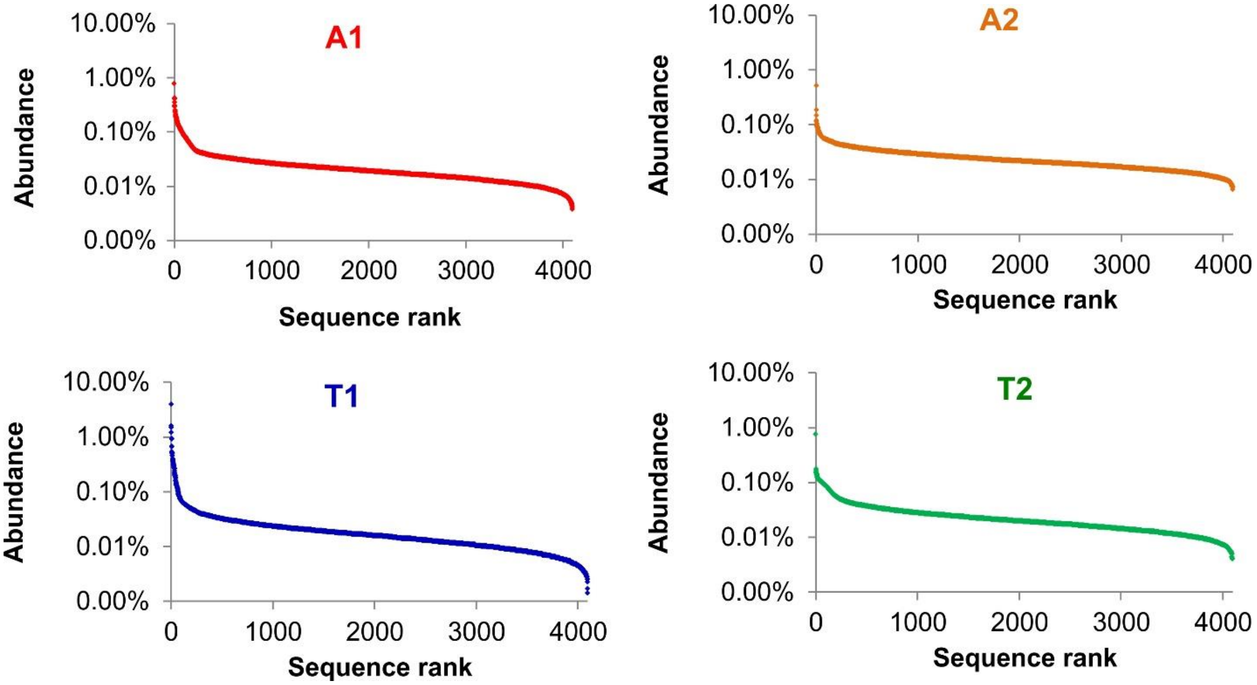

**Supplementary Figure 8.** Distribution of unique mutants in each of the four input tRNA$^{Pyl}$ libraries assessed by NGS. All possible mutants were identified in each library.

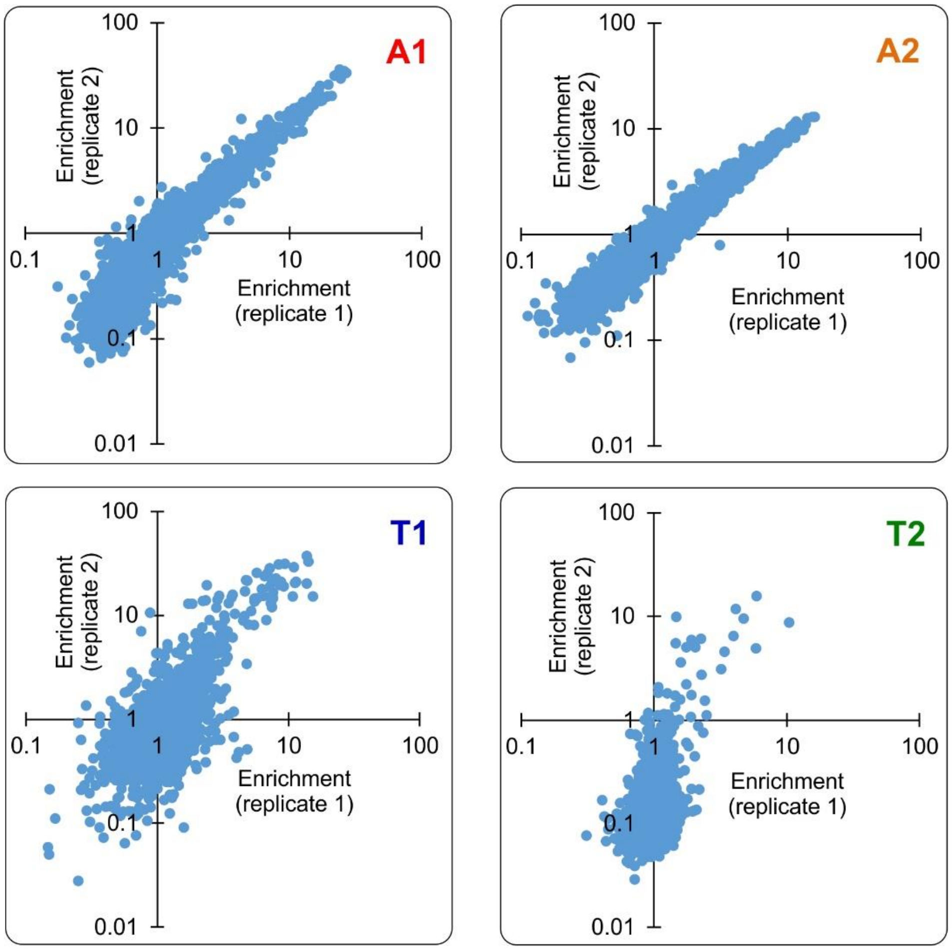

**Supplementary Figure 9.** Observed enrichment of each mutant in the four different $tRNA^{Pyl}$ libraries upon subjecting them to the VADER selection scheme. Each selection was performed in duplicate, and the normalized enrichment factors observed from each replicate were plotted against each other for each mutant. Normalized enrichments observed in duplicate selections are broadly consistent with each other. This analysis also reveals that many more mutants from A1 and A2 show robust enrichment relative to T1 and T2.

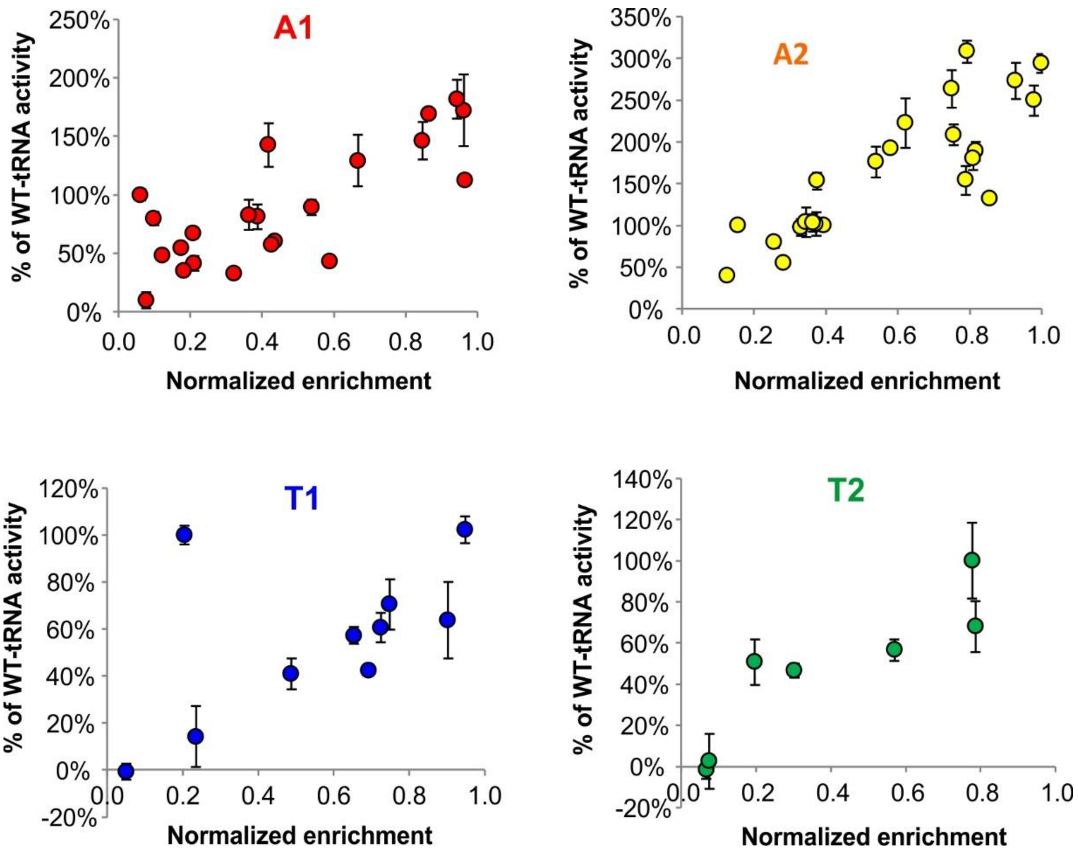

**Supplementary Figure 10.** The activity of tRNA$^{Pyl}$ mutants generally correlates with the observed degree of enrichment. The measured UAG suppression activity (Figure 2d) of individual tRNA$^{Pyl}$ mutants from various libraries are plotted against their average normalized enrichment upon VADER selection.

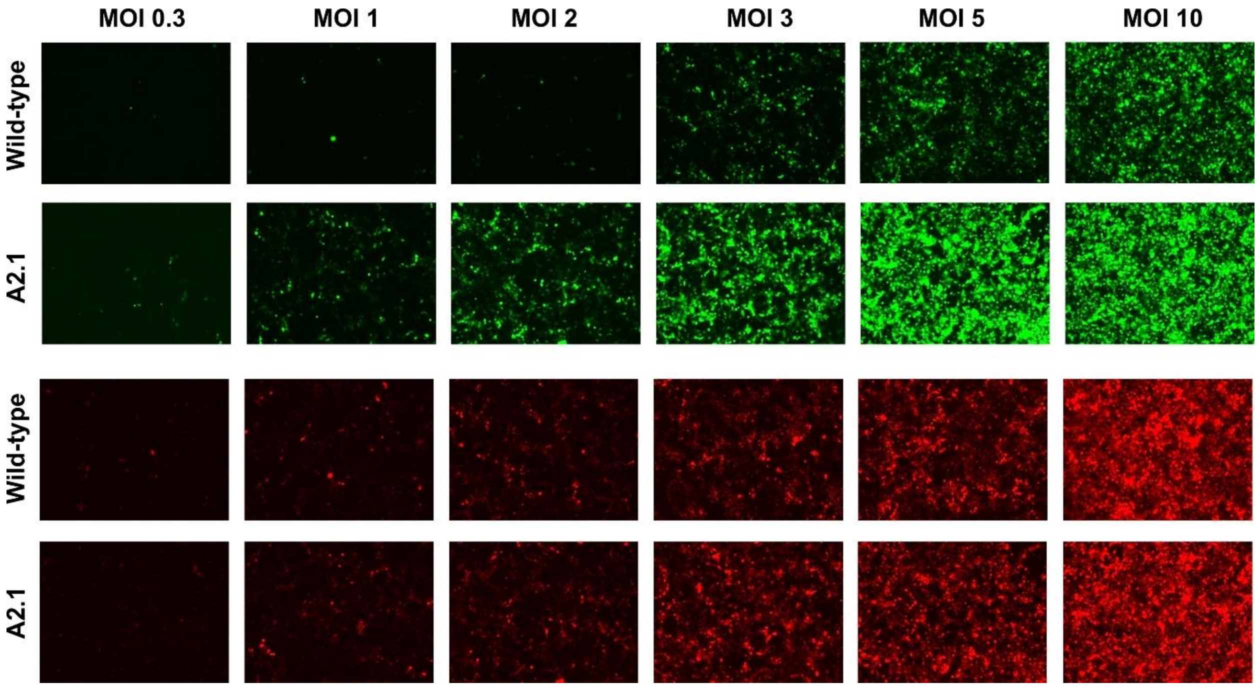

**Supplementary Figure 11**. Representative fluorescence images of cells associated with the data presented in Figure 3h and 3i. The bottom two rows show mCherry expression from increasing delivery of the tRNA-mCherry vectors. The top two rows show expression of the EGFP-39TAG reporter for the same.

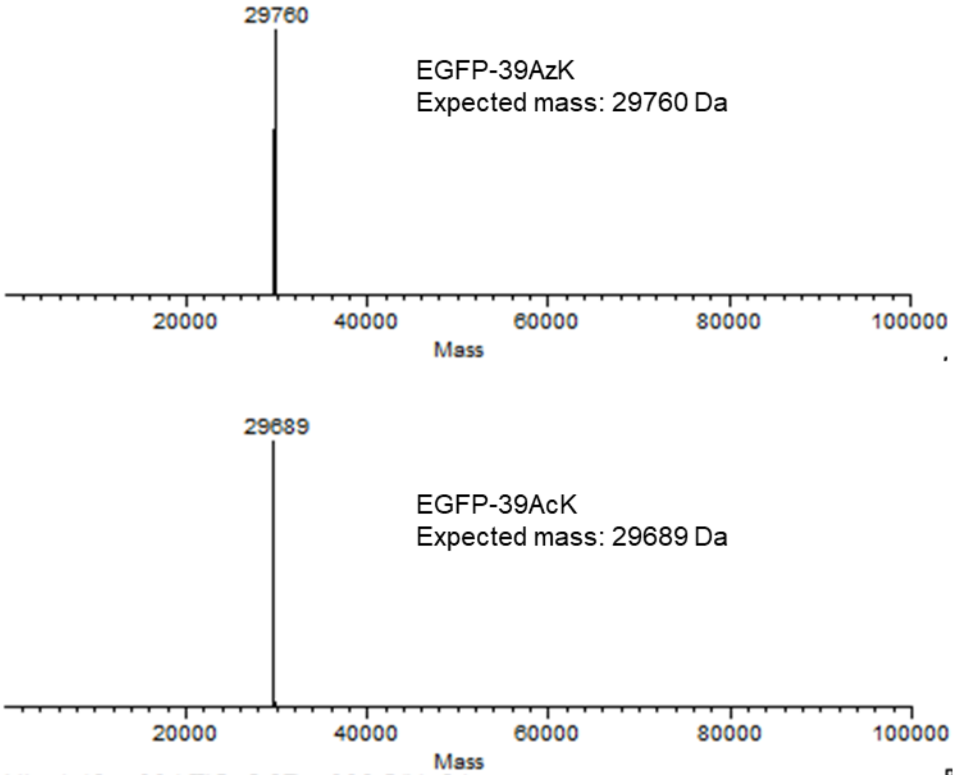

**Supplementary Figure 12**. MS analysis of purified EGFP-39TAG reporters expressed in HEK293T cells using the A2.1 $tRNA_{CUA}^{Pyl}$ variants and either wild-type MbPylRS (charging AzK) or its mutant selective for AcK, show masses consistent with the incorporation of AzK or AcK.

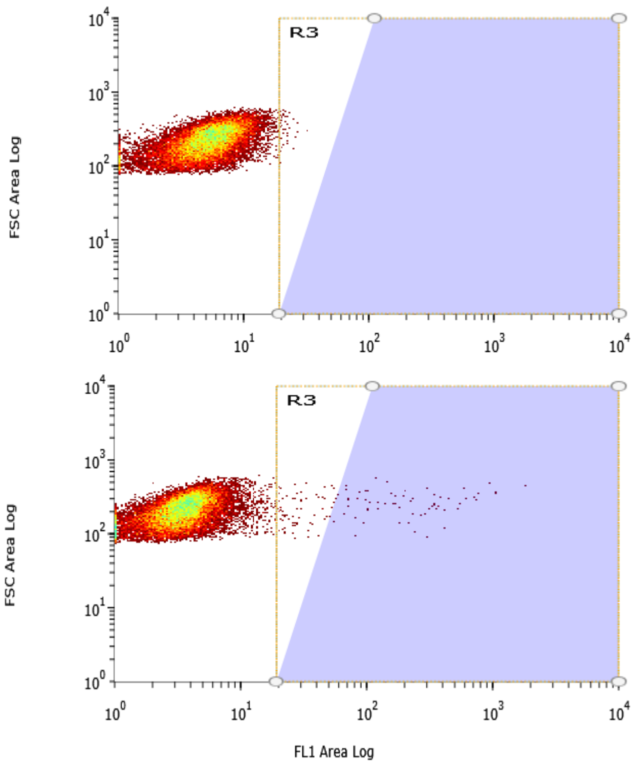

**Supplementary Figure 13.** Example of gating strategy for FACS experiments. Y-axis represent forward scatter; X-axis represent fluorescence (EGFP in this case). To titer infectivity of an AAV2 encoding the EGFP reporter HEK293T cells (12-well plate) were infected with diluted virus stock (low MOI). 48 hour later cells were detached by trypsin, washed with PBS, and analyzed by flow cytometry (bottom panel) along with uninfected HEK293T cells from the same plate as a negative control (top panel). Simple gates were drawn to exclude the population in the negative control sample.

**Supplementary Table 1**. Analysis of selected tRNA sequences (from randomly picked clones). Also see Figure 2b.

| Library | Clones sequenced | Wild-type | Non-wild-type base-paired sequences | Unique non-wild-type base-paired sequences | Not fully paired sequences |
|---|---|---|---|---|---|
| **A1** | 32 | 0 | 19 | 19 | 13 |
| **A2** | 59 | 1 | 21 | 16 | 37 |
| **T1** | 40 | 4 | 10 | 7 | 26 |
| **T2** | 35 | 8 | 7 | 5 | 20 |

**Supplementary Table 2**. Sequences of selected tRNAs that are fully base-paired (including G:U wobble pairing) from each of the four libraries. See Figure 2c for the characterization of their activity. The numbering scheme for the tRNA is shown below.

| A1 | 5 | 6 | 7 | 62 | 63 | 64 | | A2 | 2 | 3 | 4 | 65 | 66 | 67 |
|---|---|---|---|---|---|---|---|---|---|---|---|---|---|---|
| **Wild-type** | A | C | C | G | G | U | | **Wild-type** | G | A | A | U | U | C |
| **GGG/CCU** | G | G | G | C | C | U | | **GGG/CCU** | G | G | G | C | C | U |
| **GGG/CCC** | G | G | G | C | C | C | | **GGG/UCU** | G | G | G | U | C | U |
| **GGG/UCC** | G | G | G | U | C | C | | **AGC/GCU** | A | G | C | G | C | U |
| **GGC/GCC** | G | G | C | G | C | C | | **UGG/UCA** | U | G | G | U | C | A |
| **GGG/CUC** | G | G | G | C | U | C | | **AAC/GUU** | A | A | C | G | U | U |
| **GGU/ACC** | G | G | U | A | C | C | | **AUG/CAU** | A | U | G | C | A | U |
| **AGG/UCU** | A | G | G | U | C | U | | **ACU/AGU** | A | C | U | A | G | U |
| **GCU/AGC** | G | C | U | A | G | C | | **ACU/GGU** | A | C | U | G | G | U |
| **CGG/CCG** | C | G | G | C | C | G | | **UGU/ACA** | U | G | U | A | C | A |
| **CCU/GGG** | C | C | U | G | G | G | | **UGU/GCA** | U | G | U | G | C | A |
| **CCU/AGG** | C | C | U | A | G | G | | **GUG/UGC** | G | U | G | U | G | C |
| **GGA/UUC** | G | G | A | U | U | C | | **AAG/UUU** | A | A | G | U | U | U |
| **UGG/CCA** | U | G | G | C | C | A | | **GCA/UGU** | G | C | A | U | G | U |
| **GCG/CGU** | G | C | G | C | G | U | | **CAU/GUG** | C | A | U | G | U | G |
| **AAC/GUU** | A | A | C | G | U | U | | **UCG/UGG** | U | C | G | U | G | G |
| **ACA/UGU** | A | C | A | U | G | U | | **AAA/UUU** | A | A | A | U | U | U |
| **UAC/GUA** | U | A | C | G | U | A | | | | | | | | |
| **GUG/CAC** | G | U | G | C | A | C | | | | | | | | |
| **UUG/CAA** | U | U | G | C | A | A | | | | | | | | |
| **T1** | 45 | 46 | 47 | 59 | 60 | 61 | | **T2** | 48 | 49 | 55 | 56 | 57 | 58 |
| **Wild-type** | C | C | G | C | G | G | | **Wild-type** | G | G | U | U | C | C |
| **CCA/UGG** | C | C | A | U | G | G | | GG/UACC | G | G | U | A | C | C |
| **CAG/CUG** | C | A | G | C | U | G | | GG/UGCC | G | G | U | G | C | C |
| **CGG/CCG** | C | G | G | C | C | G | | AG/UGCU | A | G | U | G | C | U |
| **ACG/CGU** | A | C | G | C | G | U | | UC/AAGG | U | C | A | A | G | G |
| **CGC/GCG** | C | G | C | G | C | G | | GU/UGAU | G | U | U | G | A | U |
| **GUU/GAC** | G | U | U | G | A | C | | | | | | | | |
| **GGG/UCU** | G | G | G | U | C | U | | | | | | | | |

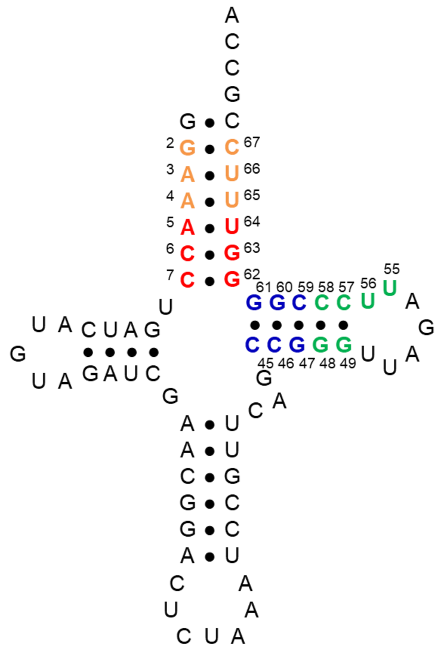

**Supplementary Table 3.** Primers used in plasmid construction, library generation, selection, and hit validation. Randomized bases (N = A, C, T, or G) are underlined.

| Primer | Sequence |
|---|---|
| **tRNA-AAV-F** | TTATTTAACGCGTCTCGCGGTCCAGTAGTGATCGACACTGC |
| **tRNA-AAV-R** | TAAATAAACGCGTGTTACGCTGGAGTCTGAGGCTCGTCCTG |
| **AccLib-short-F** | GTTTTTTGCTAGCGGATCGACGAGAGCAG |
| **AccLib-short-R** | GGTGTTTCGTCCTTTCCACAAGATATATAAAG |
| **AccLib1-NNN-F** | GATCGAACGGACTCTAAATCCGTTCAGCCGGGTTAGATTCCCGG**NNN**TTCCGTTTTTTGCTAGCGGATCGACGAGAGCAG |
| **AccLib1-NNN-R** | CCCGGCTGAACGGATTTAGAGTCCGTTCGATCTACATGATCA**NNN**TTCCGGTGTTTCGTCCTTTCCACAAGATATATAAAG |
| **AccLib2-NNN-F** | GATCGAACGGACTCTAAATCCGTTCAGCCGGGTTAGATTCCCGGGGT**NNN**CGTTTTTTGCTAGCGGATCGACGAGAGCAG |
| **AccLib2-NNN-R** | CCCGGCTGAACGGATTTAGAGTCCGTTCGATCTACATGATCAGGT**NNN**CGGTGTTTCGTCCTTTCCACAAGATATATAAAG |
| **TLib-F** | GTTTCCGTTTTTTGCTAGCGGATCGACGAGAGCAG |
| **Tlib-Short-R** | CTGAACGGATTTAGAGTCCGTTCGATCTACATG |
| **TLib1-NNN-R** | CTGCTCTCGTCGATCCGCTAGCAAAAAACGGAAACC**NNN**GGAATCTAACC**NNN**CTGAACGGATTTAGAGTCCGTTCGATCTACATG |
| **TLib2-NNN-R** | CTGCTCTCGTCGATCCGCTAGCAAAAAACGGAAACCCCG**NNNN**TCTAA**NN**CGGCTGAACGGATTTAGAGTCCGTTCGATCTACATG |
| **tRNA-Amp-F** | TTATTTACCTAGGGGTACCTCGGGCAGGAAGAGGGCCTATTTCCCATG |
| **tRNA-Amp-R** | GATGTACTGCCAAAACCGCATCACCATGGTAATAGCGATGAC |
| **PytR-evolved-TGA-F** | GGACCTGATCATGTAGATCGAACGGACTTCAAATCCGTTCAGCCGGGTTAGATTCC |
| **PytR-evolved-TAA-F** | GGACCTGATCATGTAGATCGAACGGACTTTAAATCCGTTCAGCCGGGTTAGATTCC |
| **PytR-evolved-anticodon-R** | GTCCGTTCGATCTACATGATCAGGTCC |
| **Illumina-PytR-F** | ACACTCTTTCCCTACACGACGCTCTTCCGATCTTTATATATCTTGTGGAAAGGACGAAAC |

| | |
|---|---|
| **Illumina-PytR-R** | GTGACTGGAGTTCAGACGTGTGCTCTTCCGATCGCTCTCGTCG ATCCGCTAGC |
| **Illumina-i5-D501-F** | AATGATACGGCGACCACCGAGATCTACACTATAGCCTACACTC TTTCCCTACACGACGC |
| **Illumina-i5-D502-F** | AATGATACGGCGACCACCGAGATCTACACATAGAGGCACACT CTTTCCCTACACGACGC |
| **Illumina-i5-D503-F** | AATGATACGGCGACCACCGAGATCTACACCCTATCCTACACTC TTTCCCTACACGACGC |
| **Illumina-i5-D504-F** | AATGATACGGCGACCACCGAGATCTACACGGCTCTGAACACTC TTTCCCTACACGACGC |
| **Illumina-i7-D701-R** | CAAGCAGAAGACGGCATACGAGATCGAGTAATGTGACTGGAG TTCAGACGTGTGCTC |
| **Illumina-i7-D702-R** | CAAGCAGAAGACGGCATACGAGATTCTCCGGAGTGACTGGAG TTCAGACGTGTGCTC |
| **Illumina-i7-D703-R** | CAAGCAGAAGACGGCATACGAGATAATGAGCGGTGACTGGAG TTCAGACGTGTGCTC |
| **Illumina-i7-D704-R** | CAAGCAGAAGACGGCATACGAGATGGAATCTCGTGACTGGAG TTCAGACGTGTGCTC |
| **PyltR-Tstem2-NB-R** | CCGGGAATCTAACCCGGCTGAACGGATTTAGAG |
| **5.8S-NB-R** | CGCAAGTGCGTTCGAAGTGTCGATGATCAATGTG |

**Supplementary Table 4.** Barcodes used in Illumina sequencing to enable sample multiplexing.

| Barcode | Sequence |
|---|---|
| **i5-D501** | TATAGCCT |
| **i5-D502** | ATAGAGGC |
| **i5-D503** | CCTATCCT |
| **i5-D504** | GGCTCTGA |
| **i7-D701** | CGAGTAAT |
| **i7-D702** | TCTCCGGA |
| **i7-D703** | AATGAGCG |
| **i7-D704** | GGAATCTC |

### Materials and methods

#### Cell culture.

HEK293T cells (ATCC) were maintained at 37 °C and 5% $CO_2$ in DMEM-high glucose (HyClone) supplemented with penicillin/streptomycin (HyClone, final concentration of 100 U/mL penicillin and 100 μg/mL streptomycin) and 10% fetal bovine serum (Corning). All references to DMEM below refer to the complete medium described here.

#### Statistical methods.

For assessing EGFP reporter expression in HEK293T cells, the mean of at least three independent experiments was reported, and error bars represent s.d. For evaluating the enrichment factors at each selection step, three independent experiments were performed, and for each sample three independent measurements (FACS) were made and the average of these values were reported (error represents s.d.).

#### General cloning

For all cloning, the *E. coli* TOP10 strain was used for transformation and plasmid propagation and bacteria were grown using LB for solid and liquid culture. All PCR reactions were carried out using Phusion Hot Start II DNA Polymerase (Thermo Scientific) according to the manufacturer's protocol. Restriction enzymes and T4 DNA ligase were from New England Biolabs (NEB). All DNA oligos were purchased from Integrated DNA Technologies (IDT). Sanger sequencing was performed by Eton Bioscience.

#### Unnatural amino acids

Azido-lysine (AzK) was purchased from Iris Biotech GMBH (Germany).

$N^{\varepsilon}$-acetyllysine (AcK) was purchased from Bachem.

$N^{\varepsilon}$-Boc-L-Lysine (BocK) was purchased from Chem Impex International.

#### Plasmids

The AAV2 production plasmids pHelper and pAAV-RC2 were purchased from Cell Biolabs. pIDTSmart-RC2(T454TAG)-MbPylRS has been previously described.[1] pIDTSmart-PytR plasmids with anticodons for TAG, TGA, and TAA have been previously described.[2] pAcBac1-EGFP has been previously described.[3,4]

pAAV-ITR-PytR-GFP was generated by amplifying the U6-tRNA cassette from pIDTSmart-PytR[4] with primers tRNA-AAV-F and R and inserting into the MluI site of the AAV cargo plasmid pAAV-GFP (purchased from Cell Biolabs). A KpnI site was added before the U6 promoter using primer tRNA-Amp-F. pAAV-ITR-EcYtR-GFP was constructed in a similar manner, and pAAV-ITR-PytR-mCherry was generated by replacing the GFP reporter in pAAV-ITR-PytR-GFP.

The pAAV-ITR-GFP library cloning vector was generated by cutting pAAV-ITR-PytR-GFP at two NdeI sites flanking the tRNA and ligating the resulting vector back together. This vector retains the library cloning sites but lacks a tRNA which could cause background issues during library cloning.

pIDTSmart-1xPytR-evolved was generated by amplifying the best hit tRNA (Ac2.1 GGG/CCU) from selection using primers tRNA-Amp-F and R and cloning this insert into the pIDTSmart-PytR backbone using AvrII and NheI. Anticodons were mutated using site-directed mutagenesis.

pIDTSmart-MbPylRS and pIDTSmart-AcKRS3 were generated by cloning each synthetase into the pIDTSmart backbone at the AvrII and NheI sites.

All primer sequences can be found in Supplementary Table 3.

**Library generation**

Libraries were generated by site-saturation mutagenesis using pAAV-PytR-GFP as a template. For acceptor stem libraries, the 5′ and 3′ pieces of the tRNA were first amplified using primer pairs tRNA-Amp-F + AccLib-Short-R and AccLib-Short-F + tRNA-Amp-R respectively. Randomized bases were then added by reamplifying each fragment, replacing AccLib-Short-F and R with AccLib1 or AccLib2-NNN-F and R. Finally, the fragments were joined by overlap extension PCR and then amplified using tRNA-Amp-F and R, digested with KpnI and NcoI, and ligated into the pAAV-ITR-GFP library vector which had been digested with the same enzymes. Each ligation used ~1 μg each of vector and insert. T stem library generation was similar, but for each library, only one primer containing randomized nucleotides was used. All primer sequences can be found in Table S5.

Ligations were concentrated by ethanol precipitation with yeast tRNA (Ambion) and transformed into electrocompetent TOP10 *E. coli*. >4x$10^5$ transformants were plated (>100-fold library coverage). These colonies were pooled and their DNA was miniprepped for packaging into AAV.

**Packaging and titration of mock and library tRNAs into AAV (wild-type capsid)**

To package various cargo into AAV2, 8 million HEK293T cells were seeded in a 10 cm tissue culture dish. The following day, the cells were transfected with 8 μg each of the appropriate cargo plasmid (pAAV-ITR-tRNA-fluorescent protein), pHelper, and pAAV-RC2 using polyethylenimine (PEI) (Sigma). Media was exchanged for fresh DMEM 24 hours after transfection. 72 hours after transfection, the cells were resuspended, pelleted, and lysed by freeze/thawing as previously described.[1,5] Virus was concentrated and semi-purified by PEG precipitation,[1,5] resuspended in 1 mL DMEM with FBS, and flash frozen.

#### FACS analysis to assess infective titer of AAV2 samples

Infective titers for virus preparations were determined using flow cytometry. 0.7 million HEK293T cells were seeded in each well of a 12-well plate. Next day, confluent cells (~1 million cells per well) were infected with a dilution of the AAV2 to be titered (to achieve low MOI conditions). 5 mM sodium butyrate (Sigma) was added to boost infectivity and transgene expression. Two days later, the cells were trypsinized, washed with PBS, and analyzed by flow cytometry to count the fluorescent population.

#### Transfection to determine overexpression of AAV genes with Rep and AdHelper

0.7 million HEK293T cells per well were seeded in a 12-well plate and infected the next day with AAV2 carrying a $tRNA^{Pyl}$-mCherry cargo at an MOI of 1. Cells were transfected four hours after infection with 0.6 μg pAAV-RC2 and 1.4 μg pHelper. PEI-only negative control wells received the equivalent amount of PEI to the transfected wells, but no plasmids. Virus-only wells were not transfected at all. Three days after infection and transfection, cells were lysed with CelLytic M buffer (Sigma) and mCherry fluorescence was measured on a Molecular Devices SpectraMax M5 microplate reader. The background from an uninfected well was subtracted.

#### Positive selection

8 million HEK293T cells each were seeded in three 10 cm tissue culture dishes. The next day, the cells were infected with virus containing a $tRNA^{Pyl}$ library at an apparent MOI of 5 (the actual MOI is substantially reduced in the presence of PEI, the transfection reagent). Four hours after infection, the cells were transfected with 22 μg of pHelper and 10 μg of pIDTSmart-RC2(T454TAG)-PylRS per dish using PEI. 1 mM AzK was also added at this point. One day after transfection, the culture media was exchanged with fresh DMEM containing 1 mM AzK. Cells were harvested three days after transfection and lysed as for virus isolation. The culture media was saved and recombined with clarified lysate, and this mixture was treated with 500 U universal nuclease (Thermo Scientific) for 30 minutes. Virus was recovered by PEG precipitation using 11% polyethylene glycol (Fisher) as previously described[1,5] and resuspended in 3 mL PBS.

The small-scale mock positive selections were carried out in 12-well plates. 0.7 million cells per well were seeded and infected the next day with AAV carrying a $tRNA^{Pyl}$-mCherry cargo. Four hours later the cells were transfected as described for the above selections, but with the transfection mix and AzK scaled down by a factor of 15. For PEI only wells, cells received a comparable amount of transfection reagent but no plasmid. Media was changed the day after transfection for fresh DMEM containing 1 mM AzK. Virus was harvested three days post-transfection and PEGprecipitated as described for the selections above. Confluent cells in a 12-well plate were infected with the entire output of one mock selection well and analyzed by flow cytometry.

#### Negative selection – streptavidin pulldown

The virus from positive selection (3 mL) was labeled with photocleavable DBCO-sulfo-biotin (Jena Biosciences) at a concentration of 5 μM for one hour in the dark with mixing. Immediately after labeling, excess DBCO-biotin was quenched with AzK (1 mM final concentration) and the reactions were dialyzed overnight using Slide-A-Lyzer 100 kDa MWCO devices (Thermo Scientific) against 1 L PBS at 4 °C. The dialyzed virus mixtures were split into three 2 mL tubes and each rotated overnight with 400 μL streptavidin agarose resin (Thermo Scientific) at 4 °C. The next day, each tube of beads was washed eight times with 1 mL PBS containing additional NaCl (final concentration 300 mM) with mixing between washes. Finally, the washed beads were resuspended in 8 mL PBS (300 mM NaCl) and the virus was eluted from the resin via four 30-second irradiations using a 365 nm UV diode array (Larson Electronics), with mixing between irradiations.

#### Viral DNA recovery, amplification, and cloning

The eluted virus was concentrated from 3 mL to 300 μL using Amicon Ultra-4 100 kDa MWCO centrifugal concentrators (Millipore). This mixture was heated to 100 °C for 10 minutes in order to denature the viral capsid proteins and expose the DNA. Viral DNA was then cleaned up and concentrated by ethanol precipitation using yeast tRNA (Ambion) and resuspended in a final volume of 50 μL.

20 μL of this mixture was added to a 200 μL PCR reaction and amplified with tRNA-Amp-F and R primers. The resulting DNA was digested with KpnI and NcoI and cloned into the library cloning vector using the same protocol as for original library generation.

#### Mock selections using Pyl-mCherry and Tyr-GFP

The mock selections shown in Figure 1 followed the same protocol as above, except that the starting library virus was a 1:10,000 mixture of virus made from pAAV-ITR-PytR-mCherry to virus made from pAAV-ITR-EcYtR-GFP. Mock selection results were analyzed by flow cytometry as described for virus titering, but here, cells in a 12-well plate were infected with 200 μL of the virus pool after either positive or negative selection. Red and green fluorescent cells were counted to determine the virus ratio.

#### Hit sequencing and characterization

For each library, 30-50 colonies were picked from the transformation plates generated above and sent for Sanger sequencing (Eton Bioscience). All sequences in which all randomized bases were paired were treated as potential hits, and these tRNAs were subcloned into pAAV-ITR-PytR-mCherry for further analysis.

Initial hit analysis was conducted by transfecting HEK293T cells in 24-well plates with 0.5 μg each of a potential hit pAAV-ITR-PytR-mCherry plasmid, pIDTSmart-MbPylRS, and pAcBac1-GFP(39TAG) in the presence and absence of 1 mM AzK. Two days after transfection, cells were lysed with CelLytic M buffer (Sigma) and EGFP and mCherry fluorescence were measured on a Molecular Devices SpectraMax M5 microplate reader. Values for an untransfected well were subtracted, and EGFP fluorescence was normalized to mCherry fluorescence for each well.

The best hit, A2.1 (GGG/CCU), was selected for further analysis with other stop codons and a different synthetase and Uaa, AcKRS3 and AcK. HEK293T cells in a 12-well plate were transfected with 0.375 μg pIDTSmart-PytR containing either the wild-type or evolved tRNA, 0.375 μg pIDTSmart-aaRS containing the appropriate synthetase, and 0.75 μg pAcBac1-EGFP containing one or two of the appropriate stop codon. A wild-type EGFP control well used pIDTSmart-PytR(TAG, wild-type), pIDTSmart-MbPylRS, and pAcBac1-EGFP(wild-type) in the same ratios. Two days after transfection, cells were lysed and EGFP fluorescence was measured by microplate reader. Values from an untransfected well were subtracted.

**Mass Spectrometry**

Incorporation of the correct Uaas by the evolved tRNA was confirmed by LC-ESI-MS. EGFP-39AzK-6xHis was generated by transfecting HEK293T cells in two 10 cm tissue culture dishes with 5 μg pIDTSmart-PytR-evolved, 5 μg pIDTSmart-MbPylRS, and 10 μg pAcBac1-GFP(Y39TAG) with 1 mM AzK. EGFP-39AcK-6xHis was generated by transfecting HEK293T cells in two 10 cm tissue culture dishes with 5 μg pIDTSmart-PytR-evolved, 5 μg pIDTSmart-AcKRS3, and 10 μg pAcBac1-GFP(Y39TAG) with 5 mM AcK. Cells were lysed two days after transfection using CelLytic M (Sigma), Halt protease inhibitor cocktail (Thermo Scientific), and universal nuclease (Thermo Scientific) according to the manufacturers' instructions. All proteins were isolated from clarified lysate on Ni-NTA columns using HisPur resin (Fisher) according to the manufacturer's instructions, but using 60 μL of resin and wash buffers containing 30 mM and then 40 mM imidazole. Proteins were analyzed by LC-ESI-MS using an Agilent 1260 Infinity ESI-TOF.

**Illumina sample preparation and sequencing**

To enable sequencing by the Illumina MiSeq System, DNA samples to be sequenced had Illumina adapter sequences attached via two rounds of PCR. The two consecutive reactions result in construction of the Illumina adapters provided by the TruSeq DNA HT Sample Prep Kit (Illumina). The forward adapter is

AATGATACGGCGACCACCGAGATCTACAC[i5]ACACTCTTTCCCTACACGACGCTCTTCCGATCT, wherein [i5] is an eight-nucleotide barcode sequence, and the reverse adapter is GATCGGAAGAGCACACGTCTGAACTCCAGTCAC[i7]ATCTCGTATGCCGTCTTCTGCTTG, wherein [i7] is an eight-nucleotide barcode sequence. Both i5 and i7 barcode sequences are given in Supplementary Table 4. The first set of primers, Illumina-PytR-F and Illumina-PytR-R, consist of half of the TruSeq adapters, beginning immediately after the i5 or i7 barcode, followed by primer-binding sites to anneal to the sequences surrounding the tRNA library (for the forward primer: TTATATATCTTGTGGAAAGGACGAAAC; for the reverse primer: GCTAGCGGATCGACGAGAGC). The second set of primers, a series of Illumina-i5-F and Illumina-i7-R variants containing different barcodes, consists of the 5' half of the TruSeq adapters, followed by an i5 or i7 barcode, followed by primer-binding sites to anneal to the first PCR (for the forward primer: ACACTCTTTCCCTACACGACGC; for the reverse primer: GTGACTGGAGTTCAGACGTGTGCTC).

For the first PCR, samples were prepared using the primers Illumina-PytR-F and Illumina-PytR-R and PrimeSTAR Max DNA Polymerase (Takara Bio) per manufacturer's instructions. PCR samples were purified by agarose gel extraction. A second round of PCR using Illumina-i5-F and Illumina-i7-R primer variants and PrimeSTAR Max DNA Polymerase was performed to attach the region of the adapter sequences that includes the barcodes. Unique combinations of i5 and i7 barcode sequences were applied to each sample to enable multiplex sequencing. Although both forward and reverse reads were possible, only forward reads were used in data analysis for this paper.

Samples were prepared for sequencing using the 300-cycle MiSeq Reagent Kit v2 (Illumina) per manufacturer's instructions. Sequencing was executed on an Illumina MiSeq System, with 10% Illumina PhiX Control added.

**Illumina high-throughput-sequencing data processing**

Processing of sequencing data generated by the MiSeq System was performed using bespoke Python scripts. Scripts are available upon request.

Reads were first filtered by data quality. This "Q-score filter" checked the Phred base call quality scores associated with each read in the FASTQ output files generated by the MiSeq System. Reads below a specified threshold were discarded. For this study, reads containing any bases with Q-scores lower than 14, corresponding to an error rate of 4%, were discarded. Reads were then filtered by alignment with the expected tRNA sequence. This "mismatch filter" compared each read to the expected tRNA sequence within the fixed regions of the tRNA, skipping over the randomized regions within the tRNA associated with each library. Finally, a minimum abundant library count could be specified to reduce very rare sequences that may be the result of an error. However, for these samples, the minimum abundant library count was set to 1; no library members were discarded for being too low in abundance.

For sequences that pass both the Q-score filter and the mismatch filter, the sequence regions randomized in each library were extracted and collected in a comma-separated file. For each sequence in a given sample, the "fraction of total" value was calculated by dividing the counts of a given sequence by the total counts of all sequences in that sample. The number of base pairs found within the randomized library region of each sequence was also calculated, because successful base-pairing with stem regions is important for tRNA activity.

For each library, three samples were sequenced: the input library and the output from two replicates of the selection. The counts detected for each individual sequence in each sample were tallied. For each selection, the "fold enrichment" of a given library member was determined by calculating the ratio of counts in the selection output to counts in the selection input. The "enrichment factor" of a given library member is the fold enrichment normalized to the most enriched hit, and was determined by dividing the fold enrichment of a given hit by the fold enrichment of the most enriched hit in that sample. Finally, for each library member, the average of the two enrichment factors obtained was determined, and library members were sorted by average enrichment factor.

**RNA isolation**

To generate RNA for Northern blots, 7.5 million HEK293T cells were seeded in a 10 cm tissue culture dish.  The following day, cells were transfected at 75% confluency with 24 µg of tRNA variant-containing pAAV-ITR-PytR-mCherry plasmid using polyethylenimine (PEI) (Sigma). Media was exchanged for fresh DMEM 24 h after transfection, and cells were harvested 48 h post-transfection. RNA isolation was performed with TRIzol Reagent (Thermo Fisher) following manufacturer's instructions. RNA concentration was determined via Nanodrop Spectrophotometer and integrity of the RNA was assessed by A260/A280 value, as well as the presence of distinct, intact 28S and 18S ribosomal RNA bands on 1% agarose gel.

**Northern blot probe preparation**

Oligonucleotides were 3'-end labeled with DIG using the DIG Oligonucleotide 3′-End Labeling Kit, 2nd Generation (Roche), following manufacturer's instructions. Final concentration of labelled probe was determined per manufacturer's instructions. The oligonucleotide used was PyltR-Tstem2-NB-R, which binds to the T-stem of the tRNA in order to be compatible with all A-stem variations. For detection of the 5.8S RNA positive control to assess overall RNA concentration, the oligonucleotide 5.8S-NB-R was used.

**Northern blotting**

A sensitive non-isotopic northern blot was performed using a previously established method of digoxigenin (DIG)-labeled oligonucleotide probes and 1-ethyl-3-(3-dimethylaminopropyl) carbodiimide for RNA-membrane cross-linking, with modification.[6] Denaturing gels were 8 M Urea, 6.5% acrylamide in Tris/Borate/EDTA buffer (TBE), and made in mini-gel format (10.1 × 7.3 $cm^2$ with 1.5 mm spacers). Gel wells were thoroughly rinsed with TBE after gel was set and again after pre-running gel at 250 V for 60 minutes in TBE running buffer. 5 μg of total RNA was combined with 2.5 µL Invitrogen™ Gel Loading Buffer II 2x (Denaturing PAGE, Fisher) to prepare a final sample volume of 5 µL per lane. Both RNA samples and RNA ladder (Low Range ssRNA Ladder, New England Biolabs) were denatured at 95 °C for 5 min, chilled on ice for 2 min, and then loaded onto gel. Gel was run at 4 °C for 70 minutes and then soaked in 0.05% ethidium bromide solution in RNAse free water while shaking for 10 minutes. The gel was imaged with the ChemiDoc-IT Imaging System and then soaked for 10 minutes in TBE running buffer.

A cassette sandwich was made of three sheets of 3MM Whatman paper soaked in TBE, nylon membrane, gel, and an additional three sheets of Whatman paper. Transfer of RNA from gel to membrane was done at 10 V for 60 minutes at 4 °C with the Trans-Blot SD Semi-Dry Transfer Cell (Bio-Rad).

EDC cross-linking solution was prepared as previously described.[6] Whatman paper was saturated in EDC cross-linking solution and placed on top of Saran wrap. Membrane was placed on top of Whatman paper and allowed to incubate at 60 °C for 1 hour to facilitate RNA–membrane cross-linking. Residual cross-linking solution was then removed by thoroughly rinsing the membrane with distilled water.

DIG blocking, DIG washing, DIG detection buffer were prepared per manufacturer's instructions from DIG Wash and Block Buffer Set (Millipore Sigma). CSPD detection buffer was made with 50 µL of CSPD™ Substrate (0.25 mM Ready-To-Use, ThermoFisher) and 5 mL of detection buffer. DIG antibody solution was prepared by mixing DIG antibody (Anti-Digoxigenin-AP, Fab fragments) and blocking buffer at a ratio of 1:15,000. After cross-linking, membrane was incubated in 50 mL Falcon tube at 42 °C for 30 minutes in hybridization oven with 5 mL of Invitrogen™ ULTRAhyb™ Ultrasensitive Hybridization Buffer (Fisher), preheated to 68 °C. The DIG-labelled probes PyltR-Tstem2-NB-R-DIG and 5.8S-NB-R-DIG were diluted to 25 nM and 2.5 nM, respectively, and denatured at 95 °C for 5 minutes. 5 µL of each probe was added to liquid in pre-hybridized Falcon tube and hybridized overnight at 42 °C in hybridization oven at slow rotation speed. The following day, membrane was washed through incubation twice with 5 mL Low Stringent Buffer (2× SSC with 0.1% SDS) at 42 °C for 5 minutes, twice with 5 mL High Stringent Buffer (0.1× SSC with 0.1% SDS) at 42 °C for 15 minutes, and then once with 10 mL Washing Buffer (1× SSC) at 42 °C for 10 minutes in hybridization oven. Membrane was incubated in 10 mL DIG blocking buffer at room temperature for 3 hours in hybridization oven. Membrane was then incubated with 10 mL of DIG antibody solution at room temperature for 30 minutes in hybridization oven. Membrane was then washed four times with 10 mL of DIG washing buffer for 15 minutes in hybridization oven and incubated with 5 mL of DIG detection buffer for 5 minutes at room temperature in hybridization oven. Membrane was then removed from tube with clean forceps, placed on plastic wrap, and incubated for 5 minutes with 5 mL of CSPD detection buffer.

Membrane was then placed in a heat-sealable plastic bag, sealed, and incubated at 37 °C for 15 minutes in the dark. The membrane was then imaged for chemiluminescence for 10 minutes with the ChemiDoc-IT Imaging System.

#### Production of baculovirus vectors and their use for testing tRNA$^{Pyl}$ activity with copy number control

Sf9 cells (Life Technologies) were maintained in a suspension culture at 28 ºC in Sf-900 III SFM serum-free media (Fisher Scientific). VSVG-pseudotyped baculovirus vectors were generated and titered as previously described.[3,4] For BacMam vectors containing MbPylRS and either EGFP-39TAG or wild-type EGFP, titers were determined by using the BacPAK Baculovirus Rapid Titer Kit (Clontech) per manufacturer's instructions. For tRNA-BacMam vectors, infective titers were determined using flow cytometry. 0.7 million HEK293T cells were seeded per well of a 12-well plate. The following day, confluent cells were infected with a dilution of the baculovirus suitable to achieve low MOI conditions. 5 mM sodium butyrate was added to boost infectivity and transgene expression. Two days later, the cells were trypsinized, washed with PBS, and analyzed by flow cytometry to count the population displaying mCherry fluorescence.

#### Hit characterization via baculovirus infection

0.4 million HEK293T cells were seeded per well of a 12-well plate. The following day, baculovirus was added to cells at 30% confluency. All cells received 100 MOI of the BacMam vector encoding MbPylRS and EGFP-39TAG, and 0.3-10 MOI of a BacMam vector encoding wild-type mCherry and four copies of either wild-type tRNA$^{Pyl}$ or A2.1 (Figure 3g). 1 mM BocK was also added at this time. Two days after infection, cells were lysed with CelLytic M buffer (Sigma), and EGFP and mCherry fluorescence were measured on a Molecular Devices SpectraMax M5 microplate reader. Values for an uninfected well were subtracted, and EGFP expression was normalized to the EGFP value of cells infected with 100 MOI of a BacMam vector encoding MbPylRS and wild-type EGFP.

**Data availability**. Sequences of fully base-paired selected tRNAs are available in Supplementary Table 2. Sequences of all primers used in the study are available in Supplementary Table 3. Plasmid sequences are available in Supplementary Data Set 1. NGS data sets associated with this work are available from the corresponding author upon reasonable request.
